# Human gut Actinobacteria boost drug absorption by secreting P-glycoprotein ATPase inhibitors

**DOI:** 10.1101/2022.10.13.512142

**Authors:** Than S Kyaw, Moriah Sandy, Kai Trepka, Janice JN Goh, Kristie Yu, Vincent Dimassa, Elizabeth N. Bess, Jordan E Bisanz, Peter J Turnbaugh

## Abstract

Drug efflux transporters are a major determinant of drug efficacy and toxicity. A canonical example is P-glycoprotein (P-gp), an efflux transporter that controls the intestinal absorption of diverse compounds. Despite reports that P-gp expression depends on the microbiome, the mechanisms responsible and their physiological relevance remain unclear. Surprisingly, we found that the cardiac drug-metabolizing gut Actinobacterium *Eggerthella lenta* increases drug absorption in mice through post-translational inhibition of P-gp ATPase efflux activity. P-gp inhibition is conserved in the *Eggerthellaceae* family but absent in other Actinobacteria. Comparative genomics identified genes associated with P-gp inhibition. Finally, activity-guided biochemical fractionation coupled to metabolomics identified a cluster of isoflavonoids produced by *E. lenta* related to plant-derived P-gp inhibitors. These results highlight the unexpected overlap between diet- and microbiome-derived compounds, and the importance of considering the broader relevance of the gut microbiome for drug disposition beyond first-pass metabolism.

**One Sentence Summary:** The gut bacterium *Eggerthella lenta* secretes inhibitors of P-glycoprotein ATPase activity, accelerating drug absorption.

## INTRODUCTION

Drug metabolism and disposition depend upon both genetic and environmental factors that shape intestinal metabolism, transport across the intestinal epithelium, and first-pass metabolism in the liver (Altman 2007; Brunton, Chabner, and Knollmann 2011). For the past decade, research at the interface of the microbiome and pharmacology has focused on the metabolism of drugs within the gastrointestinal tract (Zimmermann et al. 2019; Lam, Alexander, and Turnbaugh 2019; Javdan et al. 2020). While this focus is understandable given the immense and poorly characterized enzymatic diversity within the microbiome, emerging data has begun to suggest that the gut microbiome has far broader impacts on drug disposition (Bisanz et al. 2018; Kyaw and Turnbaugh 2022). Gut bacterial metabolism of common food and drug additives (*i*.*e*. excipients) increases drug bioavailability by lifting the inhibition of an intestinal drug influx transporter (Zou et al. 2020). However, it remains unclear if the diverse chemicals produced by the gut microbiome (Sugimoto et al. 2019) can also control drug influx and/or efflux transporters, and if so, whether or not these host-microbiome interactions have physiologically relevant consequences for drug bioavailability (Kyaw and Turnbaugh 2022).

Here, we begin to address this major gap in our knowledge. We opted to focus on P-gp (encoded by the *ABCB1* gene in humans and *Abcb1a* in mice) due in part to its broad relevance for >300 known endogenous and exogenous substrates, including the cardiac drug digoxin and the anti-cancer agents doxorubicin and paclitaxel (Wishart et al. 2018). *Abcb1a* can be differentially expressed between germ-free (GF) and conventionally-raised (CONV-R) mice (Hooper et al. 2001; Larsson et al. 2012); however, a recent study failed to detect a significant difference in *Abcb1a* transcript levels (Fu et al. 2017). Variation in the specific gut microbiota found between facilities could provide one potential reason for these discrepant results, consistent with experiments administering single pathogenic, probiotic, and commensal bacteria that can tune *Abcb1a* expression in either direction (Hooper et al. 2001; Saksena et al. 2011; Siccardi et al. 2008; Y. Wang et al. 2012) and a recent paper implicating butyrate-producing bacteria in colonic P-gp protein levels (Foley et al. 2021). More importantly, the functional consequences of microbiome-driven changes in *Abcb1a* expression and the mechanisms responsible remain largely unexplored.

Our prior work on the cardiac drug digoxin also motivated us to consider P-gp and its relationship to the gut microbiome. Digoxin is both a substrate for gut bacterial metabolism (Saha et al. 1983) and the model substrate for P-gp efflux (Elmeliegy et al. 2020). We previously identified a two-gene operon (the <u>c</u>ardiac <u>g</u>lycoside <u>r</u>eductase operon, *cgr*) which predicts strain-level variation in the metabolism of digoxin by the prevalent gut Actinobacterium *Eggerthella lenta* (Haiser et al. 2013). The *cgr2* gene was sufficient to catalyze digoxin reduction in a heterologous expression system and biochemical characterization of the Cgr2 enzyme suggested that its substrate scope was restricted to cardenolides (Koppel et al. 2018). Surprisingly, we also discovered that Cgr2 is necessary and sufficient for the induction of colonic T-helper 17 cells (Alexander et al. 2022; Dong et al. 2022), suggesting that dietary and/or host substrates can be reduced by this enzyme. Yet despite these mechanistic insights into the metabolic activity of *E. lenta*, the relative impacts of bacterial metabolism versus other types of host-microbiome interactions on drug disposition remain poorly understood, even for a drug as well-characterized as digoxin.

## RESULTS

### *Eggerthella lenta* increases drug absorption in mice

Given that *E. lenta* metabolizes the cardiac drug digoxin (Haiser et al. 2013), we hypothesized that mono-colonization of germ-free (GF) mice with *E. lenta* DSM2243 would significantly decrease digoxin bioavailability by enhancing first-pass metabolism. 9-week-old mixed sex GF Swiss Webster mice were mono-colonized with *E. lenta* for 4 weeks followed by oral administration of 200 μg/kg digoxin. Colonization was confirmed by qPCR (**Figure 1A**). Surprisingly, *E. lenta* increased serum digoxin levels at the first time-point (1 hour post drug administration; **Figure 1B**), resulting in a higher maximum concentration (C_max_; **Figure 1C**) and lower time to reach the maximum concentration (T_max_; **Figure 1D**) relative to GF controls. Digoxin area under the curve (AUC) was unaffected (**Figure 1E**) due to significantly decreased digoxin concentrations at 4 and 5 hours post-administration (**Figure 1B**). Taken together, these results suggest that *E. lenta* increases the rate of absorption of digoxin in gnotobiotic mice, balancing out the effects of bacterial drug metabolism (Haiser et al. 2013).

**Figure 1.**
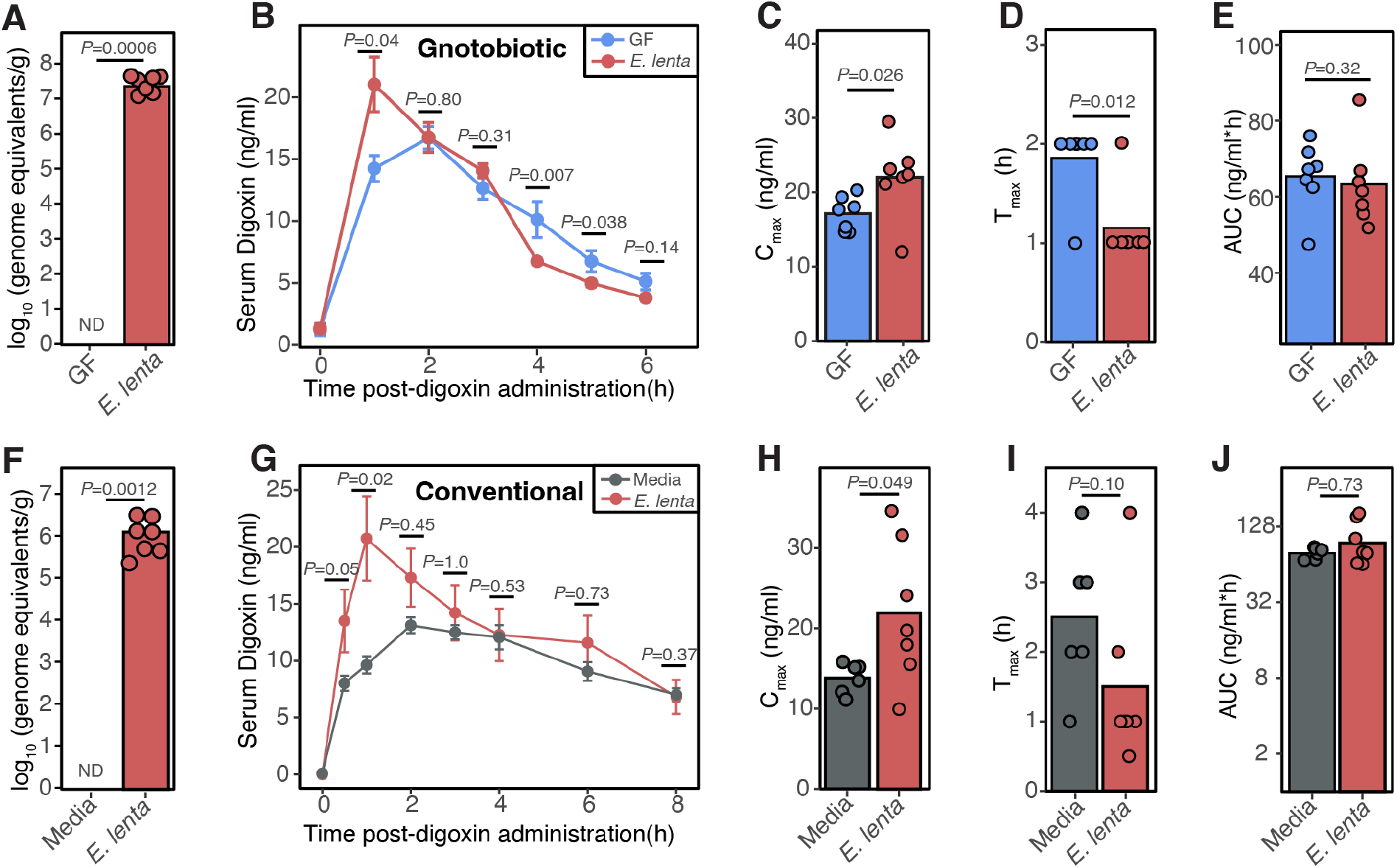
*E. lenta* increases the rate of drug absorption. **(A-E)** 9-week old GF mixed sex Swiss Webster mice were mono-colonized with *E. lenta* DSM2243 for 4 weeks followed by a single oral administration of 200 μg/kg digoxin. **(A)** Quantification of *E. lenta* DSM2243 in grams of cecal content via qPCR. No amplification was detected (ND) in the entire GF group (n=6-7 mice/group; Mann-Whitney test). **(B)** Serum digoxin was quantified at different time points in GF or *E. lenta* mono-colonized mice post-digoxin administration (n=6-7 mice/group; Wilcoxon test). From the pharmacokinetic (PK) curves, key PK parameters were quantified, such as **(C)** maximal serum concentration (C_max_), **(D)** the time at which concentrations are highest (T_max_), and **(E)** area under the curve (AUC). **(F-J)** In a separate mouse model, 8-10 week-old female CONV-R Swiss Webster mice were gavaged 10^9^ live *E. lenta* DSM2243 cells or sterile BHI^+^ media control for 7 consecutive days prior to oral administration of digoxin at 30 μg/kg. **(F)** *E. lenta* engraftment was quantified via qPCR using cecal content. BHI-gavaged group had no detectable (ND) *E. lenta* (n=6-7 mice/group; Mann-Whitney test). **(G)** Serum digoxin was quantified by LC-MS/MS at different timepoints post-digoxin administration (n=6-7 mice/group; Wilcoxon test). From the PK curves, **(H)** Cmax, **(I)** Tmax, and **(J)** AUC were quantified (Wilcoxon test).

Next, we sought to test if the observed change in digoxin absorption in response to *E. lenta* was unique to GF mice, which have broad changes to gut physiology, including changes in the expression of numerous drug transporters relative to conventionally-raised (CONV-R) specific pathogen-free animals (Fu et al. 2017; Larsson et al. 2012). To ensure consistent exposure to high levels of *E. lenta* we orally gavaged 10^9^ *E. lenta* colony-forming units (CFU) or sterile media control for 7 consecutive days to 8-10-week old female CONV-R Swiss-Webster mice prior to the oral administration digoxin. High levels of *E. lenta* were still detected in the cecum collected at the end of the experiment (**Figure 1F**). In this repeat experiment, we opted to use a lower dose of digoxin (30 μg/kg) to avoid saturating transporter effects for digoxin. The results were largely consistent with data from gnotobiotic mice (**Figures 1A-E**). *E. lenta* significantly increased serum digoxin at the 1-hour time-point following drug administration (**Figure 1G**) and significantly increased C_max_ (**Figure 1H**). There was a trend toward decreased T_max_ (*p*-value=0.10, Wilcoxon test; **Figure 1I**). Overall AUC was unchanged, due to higher levels in the control mice at the later time points (**Figure 1J**). These results demonstrate that *E. lenta* increases the rate of digoxin absorption in the presence or absence of the mouse gut microbiota.

### Human gut Actinobacterial metabolites inhibit P-gp efflux

Digoxin is a model substrate of the P-gp transporter (Elmeliegy et al. 2020), which is the only known mammalian mechanism that controls the absorption of digoxin. Drug-drug interactions resulting in altered P-gp function are known to disrupt digoxin pharmacokinetics, which can be dangerous due to its narrow therapeutic range (Elmeliegy et al. 2020). We hypothesized that *E. lenta* inhibits P-gp efflux, which would explain the elevated rate of digoxin absorption in mice (**Figure 1**). To avoid the many confounding factors in mice, we leveraged an established cell culture assay (Siccardi et al. 2008). Briefly, human enterocytes (T84 cells) were treated with Rhodamine 123 (Rh123), a fluorescent P-gp substrate, to track efflux activity under different media conditions. As expected, the P-gp inhibitors cyclosporin A (Siccardi et al. 2008), vinblastine (Siccardi et al. 2008), and verapamil (Ledwitch et al. 2016) significantly increased intracellular Rh123 levels compared to the untreated or vehicle controls (**Figure 2A**). *E. lenta* cell-free supernatant (CFS) but not cell lysates resulted in significantly increased intracellular Rh123 accumulations (**Figure 2A**). The impact of *E. lenta* CFS was dose-dependent (**Figure 2B**) and comparable in effect size to the purified pharmacological inhibitor controls (**Figure 2A**). These results suggest that one or more secreted metabolites produced by *E. lenta* inhibit P-gp.

**Figure 2.**
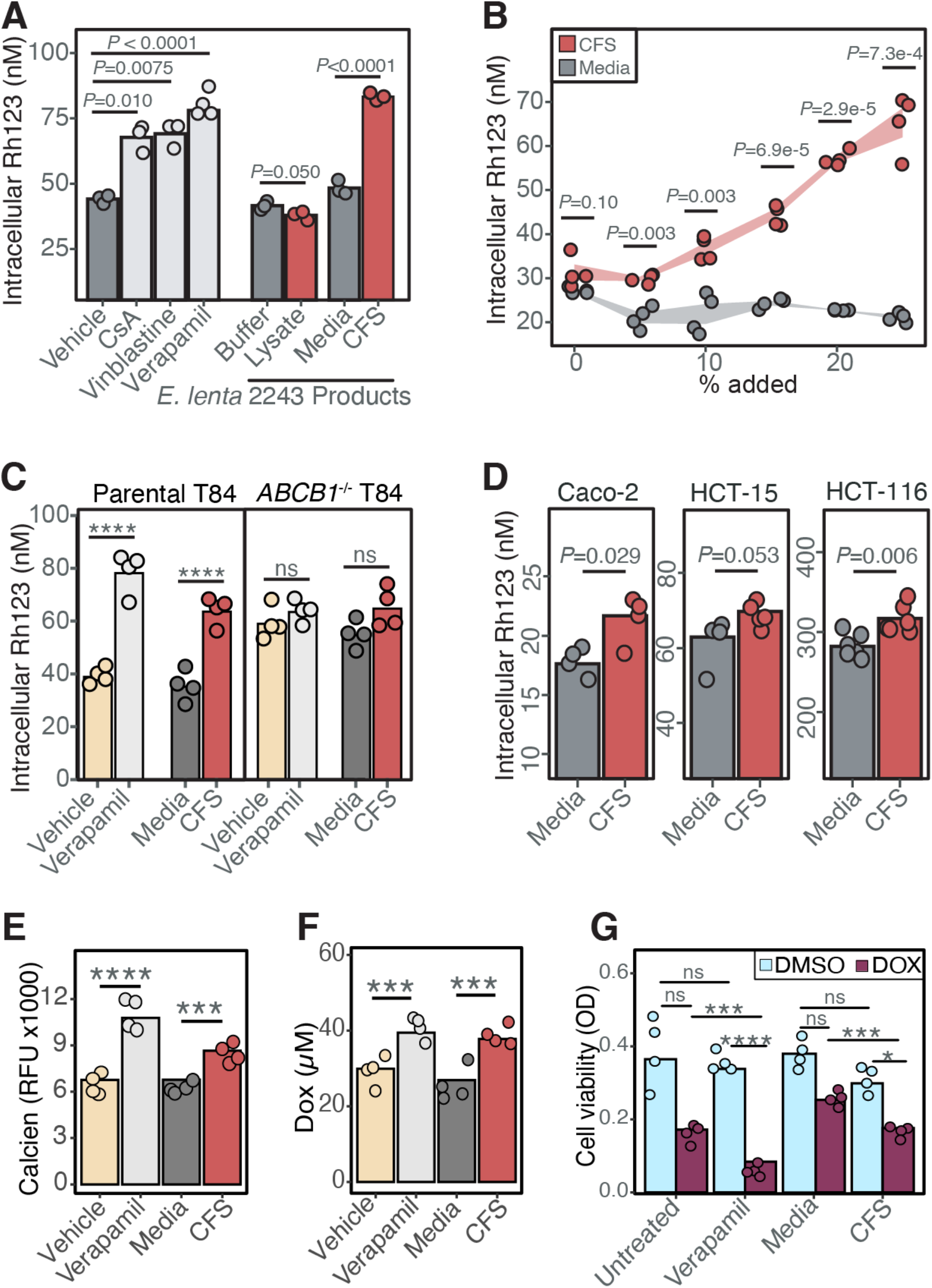
*Eggerthella lenta* inhibits P-gp efflux. **(A)** P-gp inhibition test of *E. lenta* DSM2243 cell pellet lysate and cell-free filter-sterilized supernatant (CFS) using rhodamine (Rh123) accumulation assay in T84 human enterocyte model (N=3-4/condition, ANOVA with Tukey’s post hoc test). Vehicle (DMSO), cyclosporin A (CsA), vinblastine, and verapamil were included as controls. **(B)** Dose-response test of *E. lenta* CFS on Rh123 accumulation (P_dose_=8.6e-13 ANOVA, N=4 wells/condition). **(C)** CFS Rh123 accumulation test on T84 *ABCB1*^-/-^ knockout cells compared to parental T84 cells. **(D)** CFS activity test in 3 other human enterocyte cell lines. Activity tests in T84 cells were also completed using accumulation of other P-gp substrates, such as **(E)** calcein-AM and **(F)** doxorubicin (DOX). **(G)** Synergistic effects of *E. lenta* CFS and DOX were tested using MTT viability cell assay. DMSO was used as vehicle control. Unless otherwise noted, *E. lenta* was cultured in BHI^+^. N=4/condition and 2-way ANOVA with Sidak’s multiple testing correction. *<0.05, **<0.01, ***<0.001, ****<0.0001.

We used three independent approaches to validate the inhibition of P-gp by *E. lenta* CFS: (*i*) gene deletion; (*ii*) alternative cell lines; and (*iii*) alternative substrates. *First*, an *ABCB1* knockout monoclonal cell line from parental T84 cells was generated via lentiviral delivery of *cas9* and a guide RNA against *ABCB1*. Successful gene deletion was confirmed via sequencing of *ABCB1* gene locus, Western blot, and activity testing (**Figures S1**). Similar to the verapamil positive control, incubation of *E. lenta* CFS no longer increased intracellular Rh123 levels in the *ABCB1* knockout cell line, suggesting that the mechanism is P-gp dependent and consistent with the increased basal accumulation of Rh123 in *ABCB1* knockout cells (**Figure 2C**). *Second*, incubation with *E. lenta* CFS led to high intracellular Rh123 concentrations in 3 other human enterocyte models (Caco-2, HCT-15, and HCT-116), indicating that inhibition is reproducible across cell lines (**Figure 2D**). *Third*, we validated the impact of *E. lenta* CFS on P-gp by measuring the accumulation of two additional P-gp substrates: calcein-AM (Yang et al. 2014) (**Figure 2E**) and the anti-cancer drug doxorubicin (Yang et al. 2014) (**Figure 2F**). Inhibition of P-gp efflux by *E. lenta* CFS led to a significant decrease in cancer cell viability in response to doxorubicin, consistent with the known P-gp inhibitor verapamil (Ledwitch et al. 2016) (**Figure 2G**). Taken together, these results indicate that one or more metabolites secreted by *E. lenta* robustly inhibit P-gp efflux across multiple cell lines.

### *E. lenta* inhibits P-gp ATPase activity

Small molecules and bacterial toxins can inhibit P-gp function through multiple mechanisms (**Figure 3A** and **Table S1**). Gene expression and protein levels did not explain the observed P-gp inhibition by *E. lenta. ABCB1* transcript levels were significantly increased in response to *E. lenta* CFS in 3/4 cell lines tested, with a trend towards increased expression which may indicate a compensatory mechanism (**Figure 3B**). We did not observe any changes in P-gp protein levels in response to *E. lenta* CFS (**Figures 3C**), suggesting that the major mechanism of P-gp inhibition is post-translational in nature.

**Figure 3.**
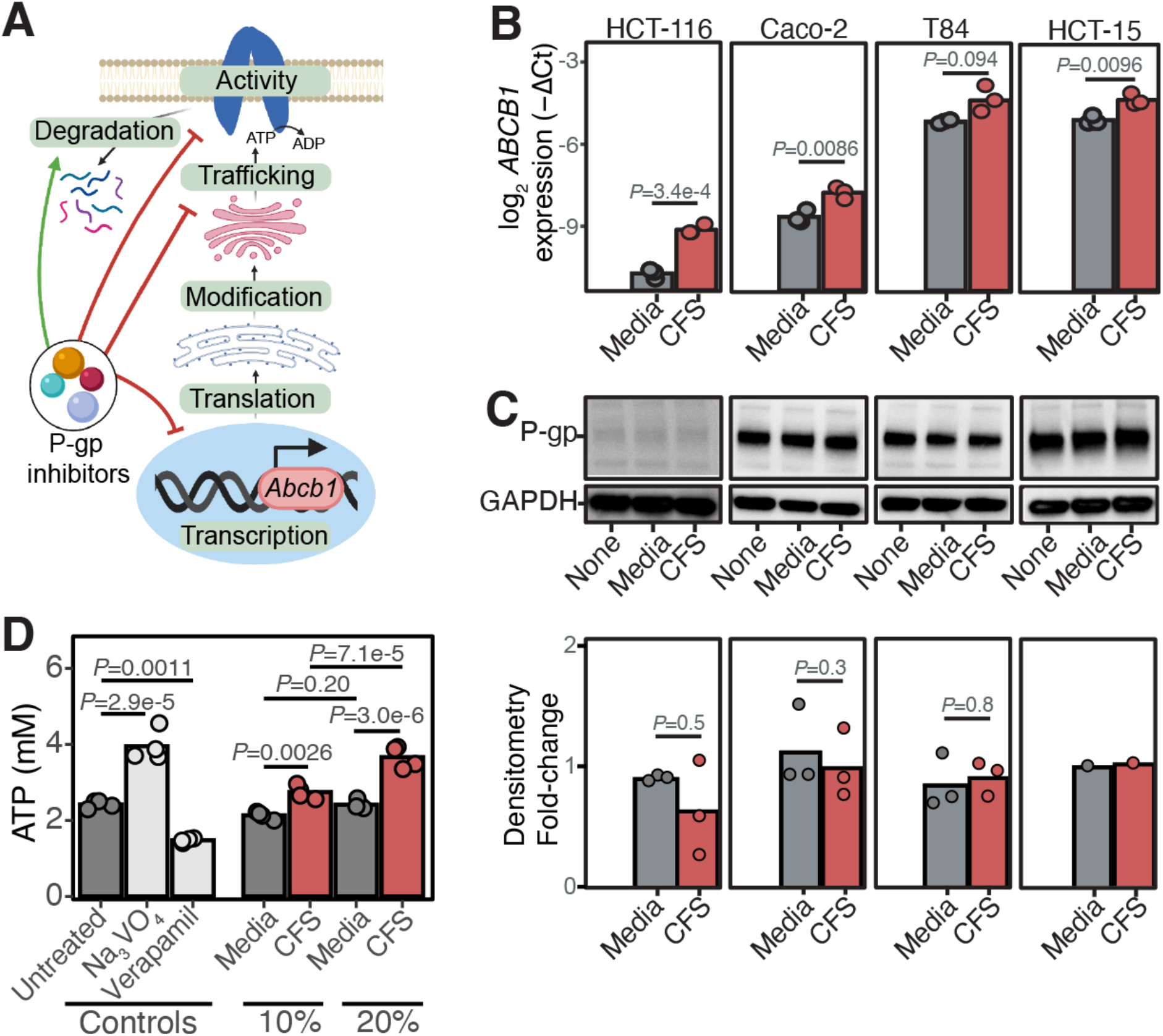
*E. lenta* inhibits P-gp ATPase activity. **(A)** Possible mechanisms of P-gp inhibition reported in the literature (**Table S1**). **(B)** RT-qPCR measurement of *ABCB1* transcription in multiple human enterocytes with varying basal expression when treated with *E. lenta* CFS. *GAPDH* and beta-actin primers were used as loading controls. Amplification cycles (Ct) were normalized to untreated conditions for each cell line (N=3 biological replicates/condition; 2-way ANOVA with Sidak’s correction). **(C)** Representative Western blots of untreated, BHI sterile media, or *E. lenta* CFS treated human enterocytes using C219 P-gp antibody and GAPDH antibody and densitometry quantifications normalized to untreated control for each cell line (N=1-3 biological replicates/condition; Wilcoxon test). **(D)** Cell-free P-gp vesicles were incubated with 3kDa filtrate of *E. lenta* CFS prepared in EDM media in the presence of ATP. At the end of the incubation period, the amount of remaining ATP was quantified. Sodium orthovanadate (Na_3_VO_4_) and verapamil P-gp inhibitors were included as controls for an ATPase inhibitor and an activator, respectively. N=4/condition; ANOVA with Tukey’s HSD post hoc test.

To test the downstream impact of *E. lenta* on P-gp activity, we turned to a cell-free vesicle system wherein the ATPase activity of P-gp can be directly quantified (Duan, Choy, and Hornicek 2009) (**Figure 3D**). The positive control (Duan, Choy, and Hornicek 2009) sodium orthovanadate significantly blocked P-gp ATPase activity whereas verapamil significantly stimulated ATPase activity (**Figure 3D**). A <3kDa fraction of small molecules from *E. lenta* CFS significantly increased free ATP in a dose-dependent manner (**Figure 3D**). These results suggest that *E. lenta* inhibits the ATPase activity of P-gp.

### Comparative genomics reveals genes associated with P-gp inhibition

Next, we sought to determine the extent to which the P-gp inhibitory trait is conserved in gut bacteria. We leveraged our previously published strain collection (Bisanz et al. 2020), testing 24 *E. lenta* strains and 10 *Coriobacteriia* relatives for their ability to inhibit P-gp, using our T84 cell-based assay. All of the tested *Eggerthellaceae* strains had a comparable degree of inhibition, on par with our positive control verapamil (**Figure 4A**). However, no inhibitory activity was observed in *Olsenella uli* (a *Coriobacteriia*) and 15 other non-*Coriobacteriia* strains, highlighting that the activity is conserved in the *Eggerthellaceae* clade but not found in more distantly-related members of the human gut microbiota (**Figure 4A**).

**Figure 4.**
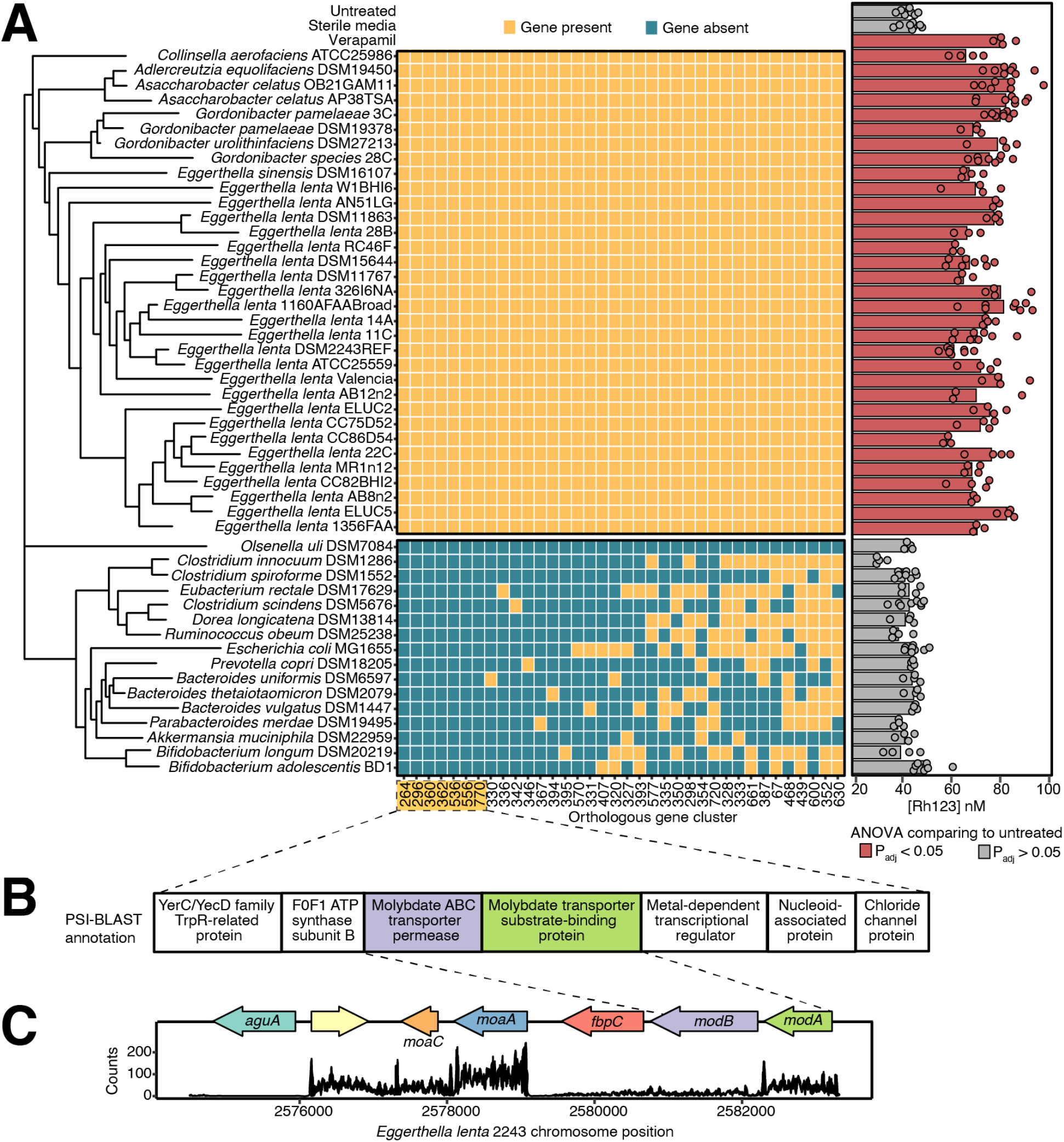
Comparative genomics reveals genes associated with P-gp inhibitory activity. **(A)** Identification of genes shared among strains with P-gp inhibitory activity assessed via Rh123 accumulation assay. Strains cultured in BHI^++^ were considered P-gp inhibitory if Rh123 accumulation is significantly higher than the untreated control. Sterile BHI^++^ media and 10μM verapamil were included as negative and positive controls, respectively (n=3-8/strain, ANOVA with Tukey’s correction). ElenMatchR and NCBI BlastX were used to identify genes unique to PGP-inhibitory bacterial strains. A cladogram shows the phylogenetic relationships between all profiled strains. **(B)** Functional annotation of 7 putative genes that were uniquely present in P-gp inhibitory strains. **(C)** Locus diagram of *modA* and *modB* conserved genes in the *E. lenta* DSM2243 genome. Counts of transcripts from *E. lenta* DSM2243 mapped to the genome are shown.

Next, we sought to use comparative genomics to determine if gene presence/absence was associated with P-gp inhibition. Using ElenMatchR, a comparative genomic tool we previously developed to identify functional genes in *E. lenta* (Bisanz et al. 2020), we identified 36 genes shared among all 33 P-gp inhibitory *Coriobacteriia* but absent in *Olsenella uli* (**Figure 4A**). From these 36 hits, 7 genes were absent from the genomes of all 15 non-inhibitory non-*Coriobacteriia* organisms (**Figure 4A; Table S2**). All 7 of these hits were transcriptionally active in *E. lenta* DSM2243 (**Table S3**). Notably, two genes predictive of P-gp inhibition were found within the same genomic locus and annotated as impacting molybdate transport: *modA* (molybdate-binding protein) and *modB* (molybdenum transport system permease protein; **Figures 4B,C**). This suggests molybdate, a redox-active metal that complexes with diverse bacterial enzymes (Zhong, Kobe, and Kappler 2020; Maini Rekdal et al. 2020), is important for the biosynthetic pathways that *Coriobacteriia* use to produce the P-gp inhibitors.

### Polar metabolites from *E. lenta* CFS inhibit P-gp

Given that the P-gp inhibitory activity was detected in the CFS (**Figure 2A**), we used activity-guided fractionation to enrich for and better understand the chemical nature of the *E. lenta* P-gp inhibitor (**Figure 5A**). To narrow down the extracellular products, *E. lenta* CFS was filtered through a 3kDa membrane. Both size fractions were tested in T84 and Caco-2 cells, which showed the activity to be in a fraction containing small molecules <3kDa in size **(Figure 5B)**. When the 3kDa filtrate was separated into polar and nonpolar fractions via liquid-liquid extraction, the activity remained in the polar fraction (**Figure 5C**). The active polar fraction was further separated via size exclusion chromatography and the activity was detected in fractions 3 and 4 (**Figure 5D**). These results indicate that *E. lenta* secretes one or more polar small molecules that inhibit P-gp.

**Figure 5.**
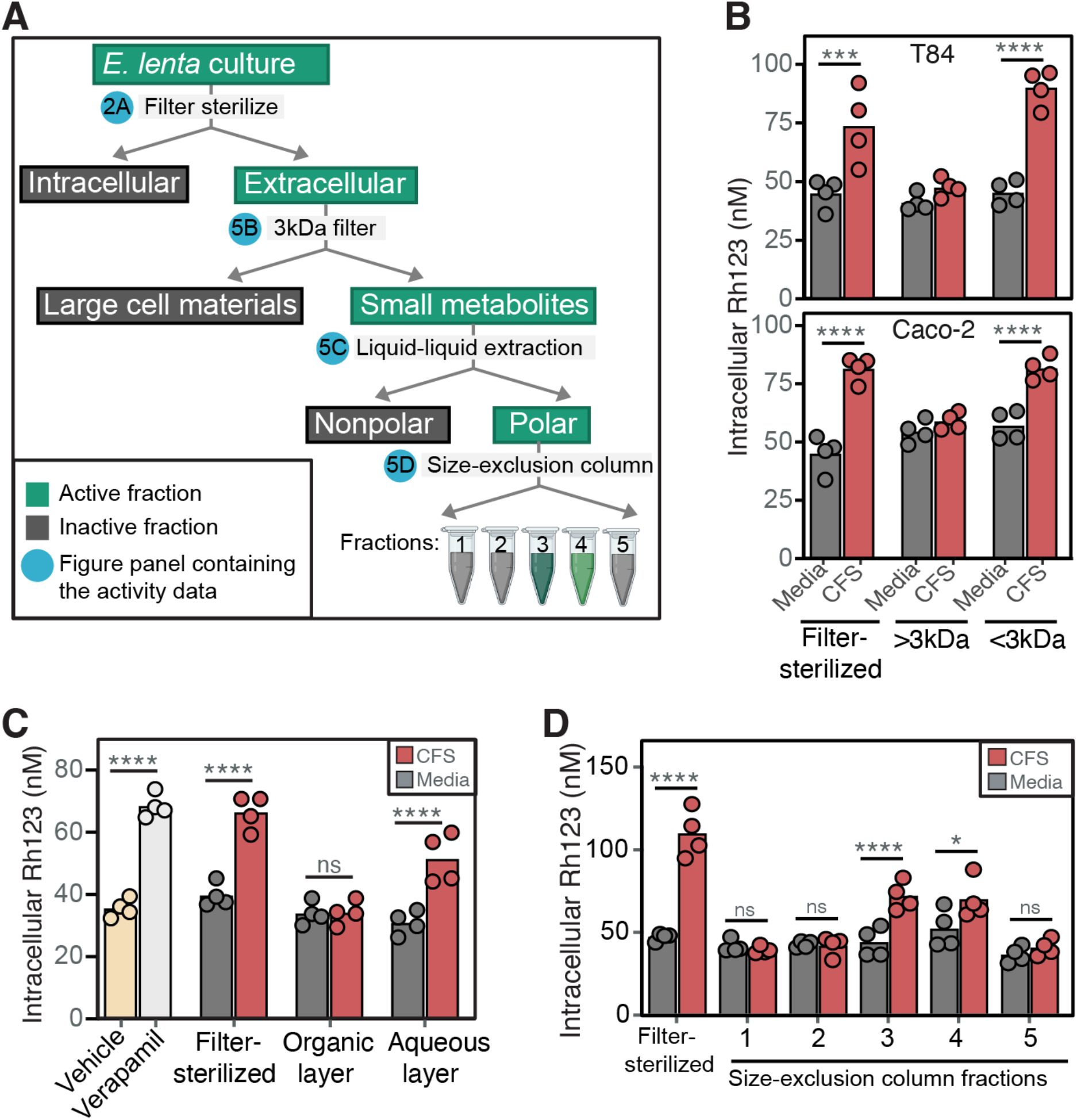
*Eggerthella lenta* secretes small polar metabolites that inhibit P-gp. **(A)** Activity-guided biochemical fractionation pipeline; the fractions are considered active when it leads to high Rh123 accumulation compared to the sterile media control. *E. lenta* CFS cultured in BHI^+^ was tested for activity in T84 cells after it was **(B)** filtered through 3kDa filter cartridge, **(C)** separated into polar and non-polar fractions via liquid-liquid extractions with MTBE:MeOH: H_2_O, and **(D)** separated into 5 fractions via size exclusion chromatography. Verapamil and filter-sterilized CFS were included as positive controls. Similar fractionations and activity testing using *E. lenta* CFS cultured in EDM are shown in **Figure S2**. N=4/condition and 2-way ANOVA with Sidak’s multiple testing correction. *<0.05, **<0.01, ***<0.001, ****<0.0001.

The arginine-supplemented BHI media used to cultivate *E. lenta* is rich and undefined, potentially complicating further efforts to identify the active compounds via mass spectrometry (Noecker et al. 2022; Bisanz et al. 2020). To address this issue, we switched to a recently developed defined minimal media that supports robust growth of *E. lenta* (EDM; **Table S4**) (Noecker et al. 2022). Consistent with our results in BHI, we noted significant P-gp inhibitory activity in *E. lenta* CFS following growth in EDM (**Figure S2**). We then repeated our activity-guided fractionation steps (**Figure 5A**) and validated the activity in our T84 assay (**Figure S2**). The biochemical fractionations not only reveal the chemical nature of the P-gp inhibitor but also enrich the metabolites that could be detected via mass spectrometry.

### Untargeted mass spectrometry reveals isoflavonoids in P-gp inhibitory fractions

Finally, we used untargeted metabolomics to identify putative P-gp inhibitors in our enriched CFS fraction in two independent experiments. In the first experiment comparing *E. lenta* DSM2243 to the EDM sterile media control, there were 25 out of 175 total features (14.3%) detected only in the active fractions (**Figure 6A, Table S5**). Molecular networking and annotations revealed that 56% of the features (14/25) unique to the active fractions were isoflavonoids (**Figure 6A; Table S5**). The remaining annotated features in the active fractions were carboxylic acids and derivatives (3/25; 12%) and benzene and substituted derivatives (3/25; 12%) (**Figure 6A**). The final 20% (5/25 features) were unannotated.

**Figure 6.**
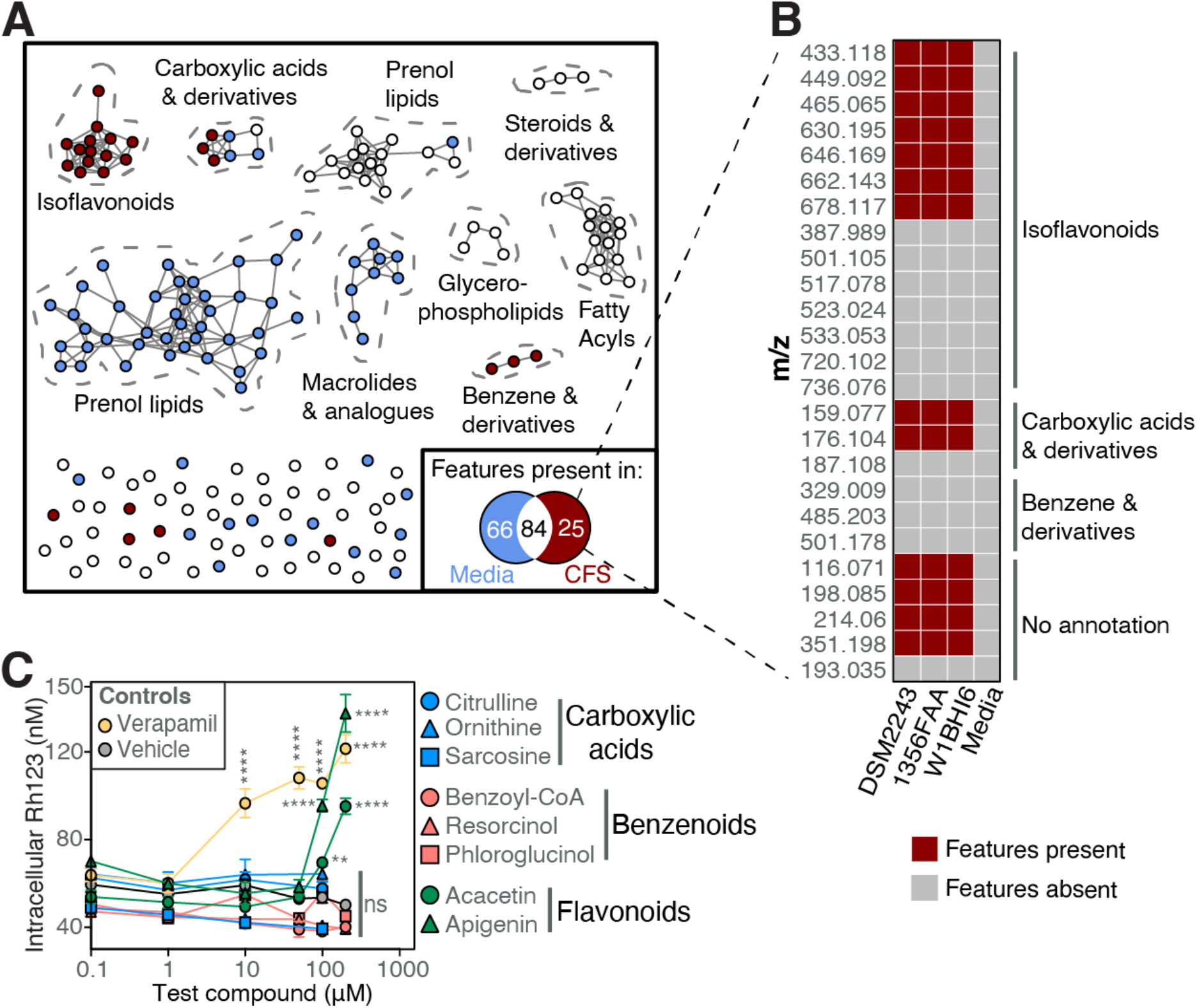
Discovery of a cluster of isoflavonoids produced by *E. lenta* linked to P-gp inhibition. Two independent untargeted metabolomics experiments were conducted (N=4/condition/experiment). In experiment 1, *E. lenta* DSM2243 CFS cultured in EDM along with sterile media control were fractionated using activity-guided fractionations (**Figure 5A**). Active fractions (**Figure S2**) were analyzed via mass spectrometry. **(A)** Feature network of features detected only in DSM2243 CFS (red points) or sterile media control (blue points), or in both (white points). Inset shows the number of unique and shared features (see **Table S5**). In experiment 2, a similar untargeted metabolomics screen was expanded to 3 *E. lenta* strains (DSM2243, 1356FAA, and W1BHI6) (**Figure S3**). 25 candidate features identified in experiment 1 were searched against strains in experiment 2. **(B)** Presence/absence heatmap of 25 candidate features in additional strains of *E. lenta*. **(C)** Dose-response curves of select carboxylic acids and derivatives, benzene and substituted derivatives, and flavonoids in T84 cells using the Rh123 accumulation assay (N=4/condition; ANOVA with Tukey’s correction). 0μM concentrations were replaced with the pseudo-value of 0.1μM during log transformation.

To test whether similar metabolites are produced in other strains with P-gp inhibitory activity, we expanded our metabolomic analysis to 3 strains in an independent experiment, including a repeat of the type strain (DSM2243). We selected *E. lenta* 1356 and *E. lenta* W1BHI6 given that they inhibit P-gp to a similar degree as *E. lenta* DSM2243 (**Figures 4A; Figure S3A**), grow robustly in the EDM media (maximum OD_600nm_: 0.8-1.0), and are furthest from DSM2243 in the phylogeny of *E. lenta* (**Figure 4A**). The EDM media is selective for *E. lenta*, prohibiting us from including non-inhibitory control strains. In this experiment, 39 out of 221 total features were detected in all 3 strains but not in the media control, including 16 isoflavonoids and 9 carboxylic acids (**Figures S3B-C; Table S6**). A combined analysis of both experiments revealed 13 features present in all active fractions but absent in all media controls: 7 isoflavonoids, 2 carboxylic acids, and 4 unannotated features (**Figure 6B**). Benzene and benzene derivatives from the first experiment were not detected in the second experiment (**Figure 6B**). Thus, these results point towards metabolites in the carboxylic acid and/or isoflavonoid groups as putative P-gp inhibitors. To further narrow down which chemical classes could explain the observed P-gp inhibition, we tested pure compounds from each group in our Rh123 assay. The carboxylic acids cluster, which contains both unique and shared features (**Figure 6A**), most likely reflects the biotransformation of arginine to other carboxylic acids. We selected the carboxylic acids citrulline, ornithine, and sarcosine for activity since these compounds are known by-products of the arginine dihydrolase pathway that *E. lenta* uses for growth (Sperry and Wilkins 1976; Noecker et al. 2022). Although benzene and derivatives were not uniquely present in CFS in the second experiment, we included benzoyl-coA, resorcinol, and phloroglucinol, which are benzene derivatives that bacteria can synthesize via the benzene degradation pathway (Harwood et al. 1998). None of the tested carboxylic acids or benzene derivatives exhibited any detectable P-gp inhibitory activity (**Figure 6C**).

Finally, we turned our attention to the isoflavonoid cluster, which represented the most common type of feature enriched in our active fraction relative to media controls. Isoflavonoids are a diverse family of natural bioactive products that have been previously shown to inhibit P-gp (Limtrakul, Khantamat, and Pintha 2005; Cui, Liu, and Chow 2019). Incubating cells with acacetin and apigenin flavonoids led to high Rh123 accumulation in our assay (**Figure 6C**), suggesting that the related isoflavonoids produced by *E. lenta* are the most likely source of P-gp inhibition.

## DISCUSSION

Our work uncovers how a prevalent gut Actinobacterium boosts absorption of digoxin, a cardiac glycoside with a narrow therapeutic window (Elmeliegy et al. 2020), by blocking a host drug efflux transporter. Mechanistic investigation revealed that *E. lenta* inhibits P-gp ATPase activity without affecting protein levels. The inhibitory activity was limited to the *Coriobacteriia* class and conserved across the *Eggerthellaceae* clade. Three distinct but complementary approaches, namely comparative genomics, activity-guided biochemical fractionations, and untargeted mass spectrometry, linked P-gp inhibition to genes involved in molybdate transport and metabolites in the isoflavonoid class.

Our results emphasize the importance of considering the vast biosynthetic potential within the human microbiome (Sugimoto et al. 2019) for explaining inter-individual variations in drug absorption and metabolism. Seminal studies have shown that microbiome-derived bioactive molecules can directly interact with human cells or receptors (host-microbe interactions) or indirectly affect the host through impacting other members of the microbiome (microbe-microbe interactions) (Sugimoto et al. 2019). One example is the production of a metabolite structurally similar to the anti-cancer drug doxorubicin, a well-known P-gp substrate (Sugimoto et al. 2019), that could compete for absorption of other P-gp substrates. As the microbiota is highly diverse and dynamic, inter-individual genetic differences in the microbiome could account for variations in drug disposition (Bisanz et al. 2018). By focusing on a prevalent member of the gut microbiota, this study demonstrates host-microbe interactions that resulted in altered drug absorption.

While bacterial metabolism of drugs is well-documented, with examples ranging from bioactivation of the antibiotic prontosil to re-activation of anti-cancer drug irinotecan (Zimmermann et al. 2019), little is known about whether and how gut microbes could alter drug absorption without biotransformation. We found that the same gut bacterial species, *E. lenta*, can have opposing effects on drug bioavailability, contributing to the intestinal metabolism of digoxin (Haiser et al. 2013; Koppel et al. 2018) while also boosting drug absorption due to inhibition of P-gp efflux. *E. lenta*, which colonizes the entire intestinal tract (Bisanz et al. 2020), could be inhibiting P-gp in the proximal small intestine, which is the major site of oral digoxin absorption (Elmeliegy et al. 2020). The remaining digoxin in the gut lumen would then be subjected to bacterial metabolism in more distal regions; however, more work is needed to map out the specific site of conversion of digoxin to dihydrodigoxin.

*E. lenta* increased digoxin absorption in both GF and CONV-R mice, which suggests that the ability to inhibit P-gp is not found in common members of the mouse gut microbiota. Interestingly, *E. lenta* is unique to humans, has a distinctive metabolic niche, and performs several unusual chemical transformations (Noecker et al. 2022), indicating that the mechanism described herein may represent a human-specific host-microbe interaction relevant to pharmacology. This host species-specificity is also true for digoxin metabolism, as the bacterial genes that metabolize digoxin to dihydrodigoxin have only been detected in select *E. lenta* strains found in human gut microbiota (Koppel et al. 2018).

Our comparative genomics results shed light on the genes necessary for P-gp inhibition by *E. lenta*. Using the clustering cutoff of protein identity <50%, we identified 7 transcriptionally-active genes that were uniquely present in the P-gp inhibitory strains. Among these 7 genes were *modA* and *modB*, two genes involved in molybdenum transport (Rech, Wolin, and Gunsalus 1996). Molybdenum, redox-active under physiological conditions, forms an active site for many bacterial enzymes involved in diverse redox chemistry (Maini Rekdal et al. 2019; Bess et al. 2020; Hille 1996; Maini Rekdal et al. 2020), potentially including isoflavonoid biosynthesis.

Although isoflavonoid biosynthetic pathways have been well-described in plants, little is known about *de novo* synthetic pathways in bacteria. Plants use phenylalanine ammonia-lyase and chalcone synthase enzymes to synthesize flavonoids from phenylalanine via the phenylpropanoid pathway (Moore et al. 2002). Once thought to be restricted to plants, homologous enzymes were found in the sediment-derived Actinobacterium *Streptomyces maritimus* (Moore et al. 2002). Biochemical studies revealed that *S. maritimus* produces flavonoids in a plant-like manner from phenylalanine (Moore et al. 2002). Although we did not detect homologues of these specific genes in *Eggerthella*, our data demonstrate that human gut Actinobacteria can also produce isoflavonoids, similar to their soil-derived counterparts. More work is needed to elucidate the biosynthetic pathway(s) responsible.

Flavonoids, typically found in fruits and vegetables, are a diverse class of natural products that exhibit numerous pharmacological properties, including anti-oxidative, anti-viral, anti-cancer, and anti-inflammatory activities (Palmeira et al. 2012). Functional assays and structure-activity relationship analyses show that several flavonoids inhibit P-gp by binding to the ATP-binding site due to their structural similarity to the adenine moiety of ATP (Palmeira et al. 2012; Fox and Bates 2007). This mechanism is congruent with our ATPase activity assay in the vesicles.

There are multiple limitations of our study. *E. lenta*-mediated P-gp inhibition was tested in GF and CONV-R mice using the P-gp substrate digoxin; synthetic communities (Cheng et al. 2022) or humanized mice (Turnbaugh et al. 2009) could be used to test how *E. lenta* affects P-gp function in the context of other members of the human gut microbiota. Although digoxin is a sensitive and physiologically-relevant P-gp probe, additional P-gp substrates could be used to assess generalizability. Our comparative genomics analysis identified 7 candidate genes that have yet to be validated. Two of the genes identified, *modA* and *modB*, share <50% sequence identity to the genes with the same annotation in *E. coli*. Since these genes have never been studied in Actinobacteria, their functions may differ from canonical *modA* and *modB* functions. We did not make genetic knockouts in *E. lenta* since the genetic tools were not available until recently (Dong et al. 2022). Through biochemical fractionations and metabolomics, we narrowed our candidate P-gp inhibitor to polar isoflavonoid compounds. Further purification efforts are required to isolate the metabolites and solve their structures, which would allow for in-depth biochemical characterization and dose response experiments in mice.

Nonetheless, our study emphasizes the broad impact of the microbiome for multiple aspects of drug disposition, including metabolism (Spanogiannopoulos et al. 2016) and absorption (Kyaw and Turnbaugh 2022; Zou et al. 2020). Continued mechanistic dissection of these and other host-microbe interactions relevant to the treatment of disease will help to explain the often unpredictable inter-individual variations in drug efficacy and toxicity and enable a new generation of microbiome-aware medicines.

## ACKNOWLEDGEMENTS

We are indebted to the other Turnbaugh lab members for their helpful suggestions during the preparation of this manuscript. We thank the Clardy lab at Harvard for their guidance in activity-guided biochemical fractionations, the Mukherjee lab at UCSF for providing *E. coli* strains with lentiviral packaging plasmids, the UCSF Gnotobiotics Core facility, the UCSF Parnassus Flow Cytometry core, and the University of Washington Mass Spectrometry Center. This project was supported by the NIH (R01HL122593, R01AT011117) and the Damon Runyon Cancer Research Foundation (DRR-42-16). JEB received fellowship support from the Natural Sciences and Engineering Research Council of Canada. TSK received support from the National Cancer Institute at the NIH (F30CA257378).

## AUTHOR CONTRIBUTIONS

Conceptualization, J.E.B. and P.J.T.; investigation, T.S.K., M.S., K.T., J.J.N.G., K.Y., V.D., E.N.B., and J.E.B.; resources, P.J.T.; writing – original draft, T.S.K. and J.E.B.; writing – review & editing, P.J.T.; supervision, J.E.B. and P.J.T.

## DECLARATION OF INTERESTS

PJT is on the scientific advisory boards for Pendulum, Seres, and SNIPRbiome; there is no direct overlap between the current study and these consulting duties. All other authors have no relevant declarations.

## INCLUSION AND DIVERSITY

We worked to ensure sex balance in the selection of non-human subjects. One or more of the authors of this paper self-identifies as an underrepresented ethnic minority in their field of research or within their geographical location. One or more of the authors of this paper self-identifies as a member of the LGBTQIA+ community. One or more of the authors of this paper received support from a program designed to increase minority representation in science. While citing references scientifically relevant for this work, we also actively worked to promote gender balance in our reference list.

## STAR METHODS

### RESOURCE AVAILABILITY

#### Lead contact

Further information and requests for resources and reagents should be directed to the Lead Contact Peter Turnbaugh.

#### Materials availability

This study does not contain newly generated materials.

#### Data and code availability

All data is available in the main text and the supplemental information and tables. Comparative genomics analyses mainly utilized the published ElenMatchR tool (Bisanz et al. 2020) and NCBI BlastX. Metabolomics datasets were analyzed on R; the script is available as a markdown HTML document on our lab GitHub (https://github.com/turnbaughlab). Any additional information required to reanalyze the data reported in this paper is available from the lead contact upon request.

### EXPERIMENTAL MODEL AND SUBJECT DETAILS

#### Experimental design

The main objective of this study was to investigate whether and how *Eggerthella lenta* alters digoxin pharmacokinetics. We utilized a combination of gnotobiotic and conventional specific-pathogen-free mouse models, cell culture assays, comparative genomics, activity-guided biochemical fractionations, and untargeted mass spectrometry, which are all outlined below.

#### Mice

All mouse experiments were approved by the University of California San Francisco Institutional Animal Care and Use Committee. The mice were housed at temperatures ranging from 19-24°F and humidity ranging from 30-70% with a 12hr/12hr light/dark cycle. No mice involved in previous procedures before experiments were used. Mice were randomly assigned to groups.

#### Gnotobiotic mouse studies

9-week-old mixed sex Swiss Webster germ-free mice were obtained from the Gnotobiotics Core Facility (gnotobiotics.ucsf.edu) at the University of California San Francisco. They were fed autoclaved chow diet (Lab Diet 5021) and were housed in gnotobiotic isolators (Class Biologically Clean) or in Iso-positive cages (Tecniplast). Mice were colonized by gavaging 200 μl of 10^9^ CFU/mL *Eggerthella lenta* DSM2243 strains and colonization was confirmed via qPCR for an *E. lenta* specific marker gene (*elenmrk1*) (Bisanz et al. 2020). They were colonized for 4 weeks prior to experiments. On the day of the pharmacokinetics experiment, pharmaceutical grade digoxin (250mcg/ml Westward) was freshly diluted in 0.3M glucose sterile solution and 200 μg/kg digoxin was orally administered to each mouse in 100-200 μl volume. 20-30 μl of blood was collected via tail vein bleeding method into a 96-well plate and allowed to clot on ice. After centrifugation at 2000g for 10 minutes at 4°C, serum was collected and stored at -80°C until mass spectrometry analysis.

#### Specific-pathogen-free mouse studies

8-10-week-old female Swiss Webster conventional mice were obtained from Taconic (Model: SW-F). They were fed a chow diet (Lab Diet 5058). Mice were orally gavaged with 200 μl of 10^9^ CFU/mL *Eggerthella lenta* DSM2243 or sterile media every day for 7 days. Daily gavage was required as *E. lenta* does not stably engraft in CONV-R mice (Alexander et al. 2022). On day 8, the digoxin pharmacokinetics experiment was conducted using the protocol outlined above by gavaging freshly prepared 30 μg/kg digoxin in 100-200 μl volume to each mouse. Similarly, serum was stored at -80°C until mass spectrometry analysis.

### METHOD DETAILS

#### Bacterial culturing

Bacterial isolates information can be found in the key resources table. All the bacterial strains were cultured at 37°C in an anaerobic chamber (Coy Laboratory Products) containing 2-5% H_2_, 20% CO_2_, and balance N_2_ gasses for 48hr. There were 3 different media used: (1) brain heart infusion (BHI; VWR 90003-040) media supplemented with 1% arginine (denoted BHI^+^), (2) BHI supplemented with 1% arginine, 0.05% L-cysteine-HCl, 5 μg/ml hemin, and 1 μg/ml vitamin K, and 0.0001% resazurin (denoted BHI^++^), and (3) *Eggerthella* selective media (denoted EDM) with media composition listed in **Table S4**.

#### Mammalian cell culture

Human colorectal cancer cell lines (T84, HCT-15, HCT-116, and Caco-2) were obtained from American Type Culture Collection and HEK-293T cell line was a gift from the Mukherjee lab. These cell lines except for Caco-2 were routinely cultured in Dulbecco’s Modified Eagle Medium/Nutrient Mixture F-12 GlutaMAX (DMEM/F12) supplemented with 10% fetal bovine serum (FBS) and 1% penicillin-streptomycin antibiotic solution. Caco-2 cells were cultured in DMEM, high glucose, pyruvate media with 10% FBS and 1% penicillin-streptomycin. Cells were grown in humidified incubators at 37°C under 5% CO_2_. Media was replaced 2-3 times a week and cells were split (1:5 for T84 and Caco2 and 1:10 for the other cell lines) using trypsin when they reached 80-90% confluence.

#### Rhodamine (Rh123) accumulation assay

Human colorectal cancer cell lines were cultured in 24-well tissue culture-treated plates. Once the cells reached 100% confluence in monolayers, they were incubated with 20% (unless otherwise noted) of CFS or sterile bacterial media control in DMEM/F12 with FBS for 18-20 hours. The media was then removed and the cells were incubated with fresh pre-warmed DMEM/F12 media containing fluorescent P-gp substrates (5μM Rhodamine 123) at 37°C under 5% CO_2_ for 30 minutes. 10μM verapamil hydrochloride, 20μM cyclosporin A, or 20μM vinblastine in 0.2% dimethyl sulfoxide (DMSO) were used as positive controls. After the media was removed, the cells were gently washed with 2mL of ice-cold phosphate buffer saline 3 times on ice. The final wash was carefully aspirated. The cells were then lysed with 300μL of RIPA buffer for 30 minutes and fluorescence was quantified using 100μL of supernatant lysate on the Biotek microplate reader (Tecan). To test P-gp inhibition with other substrates, the same procedures were performed with 50μM doxorubicin or 5μM Calcein-AM dye instead of 5μM Rhodamine 123.

#### MTT proliferation assay

T84 cells were seeded at 20,000 cells/well in 50μL in 96-well tissue culture-treated plates in DMEM/F12 supplemented with 10% FBS. To test whether *E. lenta* CFS sensitizes the cells against doxorubicin, the cells were incubated for 72 hours with 10% *E. lenta* DSM2243 CFS or sterile BHI^+^ media control and treated with either 2μM doxorubicin or vehicle DMSO control. 10μM verapamil was included as a positive control. After 3 days of incubation, the 3-(4,5-dimethylthiazol-2-yl)-2,5-diphenyl-2H-tetrazolium bromide (MTT) viability assay was performed according to manufacturer’s instructions (Promega). Briefly, 15μL MTT dye solution was added to the wells and incubated at 37°C under 5% CO_2_ for 4 hours. 100μL of solubilization solution was added and incubated for 1 hour at room temperature. Cell growth was assessed by measuring absorbance at 570nm.

#### Differential expression via qPCR

Human colorectal cancer cell lines were cultured in 24-well tissue culture-treated plates. Once the cells reached confluence, 20% cell-free supernatant (CFS) or sterile bacterial media controls were added for 18-20 hours. The media was removed and RNA was extracted using the PureLink RNA Mini extraction kit (Invitrogen), according to the manufacturer’s instructions. 5μL of 150-400 ng/μL extracted RNA was used to synthesize cDNA in 20μL reactions using iScript Reverse Transcription Supermix for RT-qPCR kit (Biorad). Diluted cDNA (1:20 in DNase-free water) was used to perform qPCR with SYBR Select Master Mix (Thermo Fisher Scientific) for 40 cycles. 30nM human *ABCB1* exon-spanning primers were used to quantify P-gp expression along with human *GAPDH* and beta-actin primers (**Table S7**) as loading controls.

#### Western Blot analysis

Human colorectal cancer cell lines were cultured in 24-well tissue culture-treated plates. Once the cells reached 100% confluence in monolayers, they were incubated with 20% CFS or sterile bacterial media control for 18-20 hours. The media was removed and the cells were washed twice with pre-warmed PBS. The cells were lysed on ice with 100μl of RIPA lysis buffer containing a cOmplete Mini Protease Inhibitor Cocktail tablet (Millipore Sigma). Proteins in the lysate were quantified using Pierce BCA Protein Assay kit (Thermo Fisher Scientific) according to the manufacturer’s instructions. After normalizing protein concentrations in RIPA buffer, 30μg of proteins in Laemmli sample buffer were loaded in a 10-well 4-15% Mini-PROTEAN gradient SDS-PAGE gel (Biorad) and ran for 15 minutes at 70V followed by 150V for 45 minutes in Tris-Glycine-SDS buffer at room temperature. Precision Plus Protein Dual Color standards were included in each gel. The proteins in the gel were then transferred to polyvinylidene difluoride membranes overnight at 23V at 4°C in the N-cyclohexyl-3-aminopropanesulfonic acid buffer. After blocking membranes in 5% milk, the membranes were incubated with C219 anti-ABCB1 primary antibody (Genetex) at 1:200 or anti-GAPDH primary antibody (Invitrogen) at 1:5000 in 5% milk in PBS-T blocking buffer overnight at 4°C. The membranes were washed 3 times and incubated with goat anti-mouse HRP-conjugated secondary antibody (Abcam) at 1:10,000 in a blocking buffer for 1 hour at room temperature. After washing the membranes, they were treated with Clarity Max Western ECL substrates (Biorad) for 5 minutes and visualized in the ChemiDoc (Biorad). The densitometry analysis was performed using the Biorad ImageLab software.

#### P-gp vesicle assay

To test whether *E. lenta* modulates P-gp ATPase activity, vesicles containing P-gp (5mg/ml) purchased from Thermo Fisher were tested for ATP consumption in P-gp Glo assay (Promega) according to the manufacturer’s instructions. Briefly, the vesicles were thawed rapidly at 37°C and immediately kept on ice. The vesicles were diluted to 1.25mg/ml with Pgp-Glo assay buffer before each assay. For each assay 20μl of 1.25mg/ml vesicles was mixed with 10-20μl of test agents (CFS, sterile media, 0.5mM verapamil, 0.25mM Na_3_VO_4_, or Pgp-Glo assay buffer control) in the presence of 10μl of 25mM MgATP in white flat-bottom 96-well plates. Pgp-Glo assay buffer was added if necessary to achieve the total volume of 50μl. The reaction was incubated for 1 hour at 37°C under 5% CO_2_.

After incubation, the reaction was diluted 1:4 (12.5μl into 37.5μl of P-gp-Glo assay buffer) in a new white flat-bottom 96-well plate and 50μl of ATP detection reagents were added. Standard curves (3mM - 0.375mM MgATP at 1:2 dilution) in different matrices (CFS, sterile media, Pgp-Glo assay buffer) were included. The mixture was incubated at room temperature for 20 minutes and luminescence was quantified on a plate reader (Tecan). Luminescence measurements were converted to ATP concentrations using matrix-specific standard curves.

#### Generation of P-gp knockout monoclonal cell line

*Escherichia coli* strains containing pCMV-dR8.91 and pMD2.G lentiviral packaging plasmids were a kind gift from the Mukherjee lab. An *E. coli* strain carrying the transfer plasmid (pCas9) with cas9 gene, eGFP, neomycin, and ampicillin marker genes and a separate *E. coli* strain carrying the transfer plasmid (pABCB1) with sgRNA against human *ABCB1* gene, mCherry, puromycin, and ampicillin maker genes were purchased from Genecopoeia. To harvest the plasmids, these strains were cultured separately in LB media supplemented with 50μg/ml carbenicillin aerobically overnight with shaking at 37°C. The resulting turbid cultures were centrifuged to collect the cell pellets, which were then processed with Midi-Prep plasmid extraction kit (Qiagen) based on the manufacturer’s protocol. Plasmids were confirmed via enzymatic restriction digest.

4μg of pCMV-dR8.91, 1μg of pMD2.G, and 5μg of either pCas9 or pABCB1 plasmids were added to a tube containing 30μL of LT1 transfection reagent (Mirus) and 1mL of Opti-MEM I Reduced Serum medium. The contents were vortexed and incubated at room temperature for 30 minutes. The mixture was added drop-by-drop into a 10cm cell culture dish containing HEK293T cells at 90% confluency. The cells were incubated for 48 hours and the supernatant containing recombinant lentiviral particles was filtered using 0.45μm filters. The lentiviral filtrate was flash frozen.

To transduce the cells, 0.5ml of lentiviral supernatant and 0.5ml of 20μg/ml polybrene transfection agent in cell culture media were added to T84 cells cultured in 12-well plates at 70% confluency. After 72hr of incubation, 10μg/ml of puromycin and 3mg/ml of geneticin antibiotics were added to select for successfully transduced cells. After 10 days of antibiotics selection, GFP^+^/RFP^+^ double-positive cells from the resulting polyclonal pool were sorted individually into 96-well plates using a cell sorter at the Flow Cytometry Core. The sorted cells were expanded into monoclonal cell lines, which were tested for *ABCB1* deletion via PCR, Western blot, and functional rhodamine accumulation assay.

#### Comparative genomics and associated transcriptomics

Comparative genomics among *Coriobacteriia* was performed using ElenMatchR (Bisanz et al. 2020). Genes were clustered at a minimum of 50% amino acid identity and 80% query coverage to allow for identification of orthologs across genus boundaries. Hits from ElenMatchR were compared to the genomes of non-Coriobacteriia non-inhibitory strains using NCBI BlastX with identical cutoffs for amino acid identity and query coverage and a liberal E value threshold of 10 to ensure that all identified gene clusters were unique to P-gp inhibitory bacterial strains. Gene annotations were called using *E. lenta* DSM2243 genes corresponding to each orthologous gene cluster hit. Raw *E. lenta* DSM2243 RNA sequencing data in BHI was obtained from and filtered and aligned as described (Maini Rekdal et al. 2019). Genes with RPKM >10 in all 3 replicates were called transcriptionally active.

#### Activity-guided biochemical fractionations

*E. lenta* DSM2243 cultures were grown anaerobically at 37°C in BHI^+^ or EDM media with sterile media control for 48 hours. To test whether *E. lenta-derived* P-gp inhibitor was intracellular or extracellular, cell pellets and supernatants were separated via centrifugation to test for P-gp inhibitory activity using our Rh123 assay. The pellet was reconstituted in 15 mM HEPES buffer (pH 7.3), lysed using bead beating with Lysing Matrix E tubes (MP Bio) in a Mini Beadbeater 16 Disrupter 16 (BioSpec Products), and tested for activity. Supernatant was pH-adjusted to 7.6 with NaOH or HCl, filter-sterilized using 0.22 μm filters to create CFS, and tested for activity. To test whether the inhibitor was large cellular materials or small metabolites, CFS was further separated using 3kDa filter cartridges. Large materials on the filter membrane were reconstituted in sterile media in the same volume. The flowthrough and reconstituted large materials were then tested for activity. To further separate the small metabolites based on polarity, 4ml of 3kDa flowthrough was added to 12ml of M1 solvent (75% MTBE and 25% MeOH) and 8ml of M2 solvent (25% MeOH in water), vortexed for 1 minute, and centrifuged at maximum speed for 5 minutes at 4°C. The polar aqueous layer was lyophilized down to 2ml while the non-polar organic layer was dried under a gentle stream of nitrogen gas in TurboVap. Their activity was tested after reconstituting in 0.5x volume of water to account for incomplete extractions and compound loss through prior steps. To fractionate based on size, 1ml of active polar fraction was loaded to G10 size exclusion midi-columns. Once the solution was absorbed into the column, water was added on top gently without disturbing the beads and allowed to flow through via gravity. 1ml fractions were collected and tested for activity.

#### Untargeted metabolomics

Untargeted liquid chromatography high-resolution mass spectrometry (LC-HRMS) analysis was performed on a Sciex Exion UPLC equipped with a BDS C8 hypersil column (Thermo, 150 × 4.6 mm, particle size 5μm) and coupled to a Sciex TripleTOF 6600+ operated in positive mode electrospray ionization with IDA acquisition of MS2 (accumulation time 0.25 s, collision energy 35V (15V spread), acquisition window 100-2000 Da). A gradient of 0-100% acetonitrile over 14 min was used for compound separation.

Feature-based molecular networking analysis (FBMN) (Nothias et al. 2020) was performed on shared ms/ms fragments across all metabolite features to determine the relationship between metabolite features detected in our HPLC fraction mixtures. Raw mass spectrometry data were preprocessed using MzMine2 (Pluskal et al. 2010; Katajamaa, Miettinen, and Oresic 2006) for feature detection, chromatogram building, peak deconvolution, isotope grouping, gap filling, and alignment. The resulting intensity-based metabolite feature matrix and MS2 mgf files were used for further statistical and network analyses in Metaboanalyst (Pang et al. 2021) and GNPS (M. Wang et al. 2016). Following FBMN, Network Annotation Propagation (NAP) (da Silva et al. 2018), MS2LDA (Wandy et al. 2018), Dereplicator (Mohimani et al. 2018), and MolNetEnhancer (Ernst et al. 2019) were used to assign chemical classifications to the metabolite features. To summarize, a feature-based molecular network was created using the online workflow (https://ccms-ucsd.github.io/GNPSDocumentation/) on the GNPS website (http://gnps.ucsd.edu). MS/MS spectra were window filtered by choosing only the top 6 fragment ions in the +/- 50Da window throughout the spectrum. The precursor ion mass tolerance was set to 2.0 Da and an MS/MS fragment ion tolerance of 0.5 Da. A network was then created where edges were filtered to have a cosine score above 0.7 and more than 6 matched peaks. Further, edges between two nodes were kept in the network only if each of the nodes appeared in each other’s respective top 10 most similar nodes. Finally, the maximum size of a molecular family was set to 100, and the lowest scoring edges were removed from molecular families until the molecular family size was below this threshold. The spectra in the network were then searched against GNPS’ spectral libraries. The library spectra were filtered in the same manner as the input data. All matches kept between network spectra and library spectra were required to have a score above 0.7 and at least 6 matched peaks. The molecular networks were visualized in R.

#### Quantification of serum digoxin

To quantify serum digoxin from the pharmacokinetics experiments, 5μl serum was combined with 100μl of 10ng/ml oleandrin internal standard and 5μl of MeOH. For controls, 5μl of 0-200ng/ml (1:2 dilution series) digoxin/dihydrodigoxin mix in MeOH was added to 5μl of blank serum along with 100μl of 10ng/ml oleandrin. To all tubes, 1ml of 1:1 MTBE:ethyl acetate was added and vortexed for 2 minutes. After centrifuging for 10 minutes at 850 xg, the samples were dipped into the dry ice/isopropanol bath to freeze the aqueous layer. The liquid organic layer was transferred into new 2ml tubes and dried to completion under a gentle stream of nitrogen gas at 10psi at room temperature in a turbovap. The dried samples were reconstituted in 100μl of 65% MeOH in water and stored at -20°C. The extracted samples were analyzed in two methods.

In method A, electrospray ionization (ESI) triple-quadrupole liquid chromatography-mass spectrometry in negative-ion mode with multiple reaction monitoring was used to detect digoxin, dihydrodigoxin, and oleandrin. 10μl of the samples were injected onto a column (ODS Hypersil 4.6 × 100 mm, 5 μm particle size) using Shimadzu SIL-30AC injector set at 4°C. The column temperature was maintained at 50°C. Using 65% MeOH in water as a mobile phase in isocratic flow, each injection was run for 5 minutes. Digoxin, dihydrodigoxin, and oleandrin were expected to elute at 2.0 minute, 2.1 minute, and 3.6 minute, respectively. Expected Q1 - Q3 m/z masses were as follows: 779.4 - 475.2 for digoxin, 781.4 - 521.1 for dihydrodigoxin, and 575.2 - 531.1 for oleandrin. Peak areas of the standard curves were converted to concentrations at ng/ml using weighted linear regression (1/x^2^).

In method B, the extracted samples were shipped on dry ice to the University of Washington Mass Spectrometry Center. Waters Xevo XS tandem mass spectrometry was used to detect digoxin, dihydrodigoxin, and oleandrin. 10μl of the samples were injected onto a column (Waters BEH C18, 2.1×50mm, 1.9μm particle size) and ran for 7 minutes. Gradient mobile phase was used (A: 0.1% formic acid in H_2_O and B: 0.1% formic acid in MeOH) at 50:50 for 0-4 minutes, 20:80 for 4-5 minutes, and 50:50 for 5.1-7 minute at 0.3ml/min. A post-column infusion of 5mM ammonium acetate was used at 10μl/min. The expected retention times for digoxin, dihydrodigoxin, and oleandrin are 2.75min, 2.6min, and 3.41min respectively. NH_4_-adducts for digoxin (parent m/z: 798.6; daughter m/z: 651.6) and dihydrodigoxin (parent m/z: 800.6; daughter m/z: 408.3) were monitored. Oleandrin was monitored at 577.4 m/z for the parent compound and 373.2 m/z for the daughter compound. Peak areas of the standard curves were converted to concentrations at ng/ml using linear regression.

#### Quantification of *E. lenta* in mice

0.1-0.6g of cecal content collected from mice at the end of the pharmacokinetics experiments was used to extract DNA using the ZymoBIOMICS DNA extraction kit according to the manufacturer’s instructions. 2μL of extracted DNA, 2μL of water, 5μL of iTaq Supermix reagents, and 1μL of 2μM *E. lenta*-specific primers with a FAM fluorophore probe (*elnmrk1*; see **Table S7**) were amplified in 4 technical replicates using the following parameters for 40 cycles: initial denaturation at 95°C for 5 minutes, denaturation at 95°C for 5 seconds, and anneal/extension at 60°C for 15 seconds. A 10-fold dilution series of DNA from *E. lenta* DSM2243 culture was included to construct a standard curve to convert amplification cycle (Cq) to genome equivalents using the formula that 1ng of DSM2243 DNA = 2.7e5 genome equivalents.

### QUANTIFICATION AND STATISTICAL ANALYSIS

All statistical analyses and data visualizations were performed in R v4.1.1 or PRISM 9.4.1. Statistical tests, sample size, standard error, and multiple testing correction methods are reported in the figures and figure legends.

## SUPPLEMENTARY FIGURES AND LEGENDS

**Figure S1.**
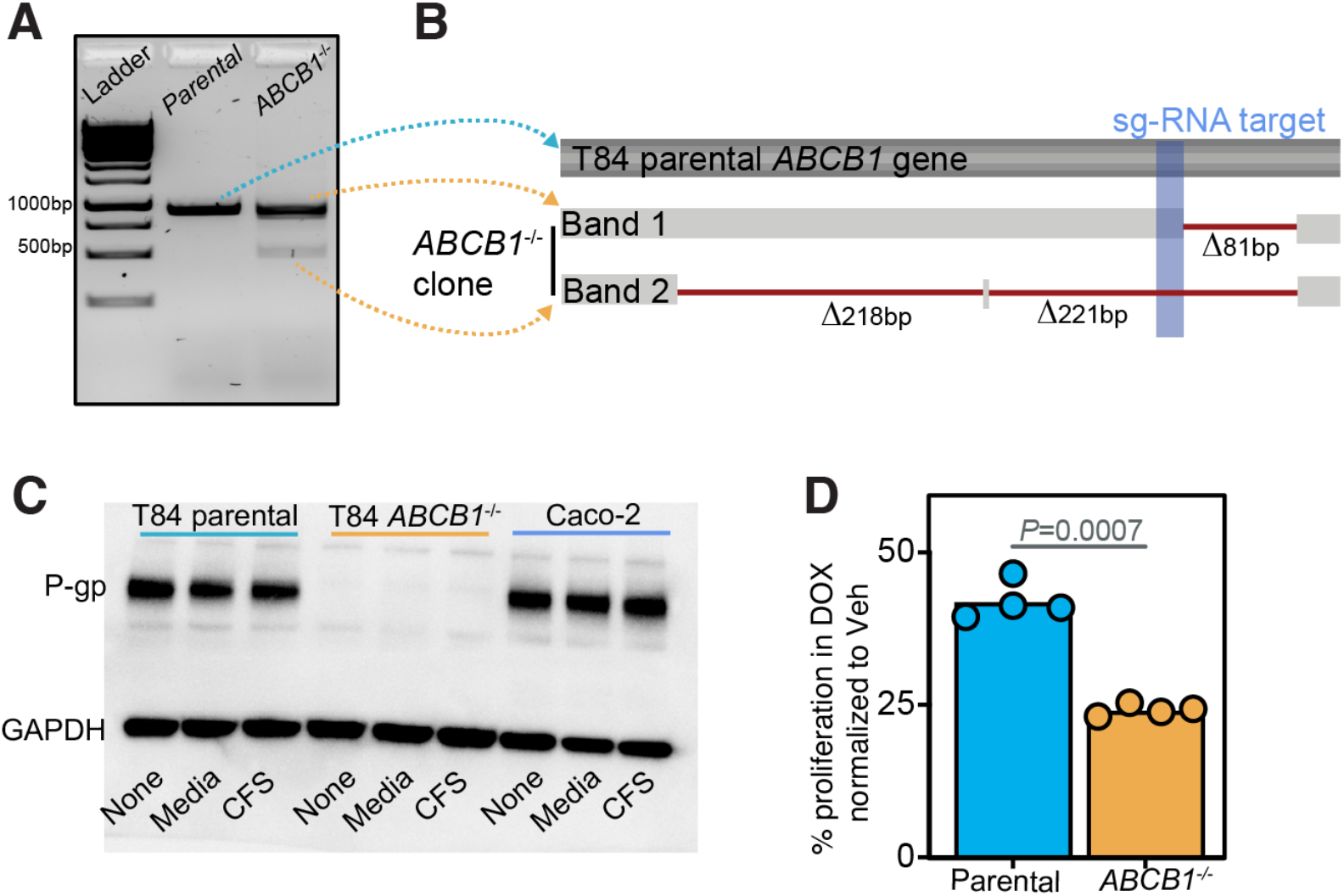
related to Figure 2. P-gp knockout monoclonal cell line generated from T84 parental cells is confirmed via sequencing, Western blot, and activity assay. **(A)** T84 cells were transduced with lentiviral particles carrying cas9 gene and small guide RNA (sg-RNA) against *ABCB1* gene. The transfected cells were sorted into a single cell per well and expanded to generate a monoclonal cell line. DNA was extracted and PCR amplified using primers flanking the sg-RNA target site. **(B)** The two visible bands in the gel were extracted, sequenced, and aligned to the parental T84 PCR product using NCBI multiple alignments tool. **(C)** Western blot of untreated (none), BHI^+^ media, or *E. lenta* CFS treated *ABCB1*^-/-^ cell lysates. T84 parental cells and Caco-2 cells were included as controls. **(D)** P-gp effluxes doxorubicin (DOX) rendering cells with functional P-gp more resistant. Cells without functional P-gp are therefore more sensitive to DOX. Using these principles, P-gp function in *ABCB1*^-/-^ cells was tested in a DOX sensitivity assay and growth was normalized to DMSO vehicle (Veh) control (N=4/condition, Wilcoxon test).

**Figure S2.**
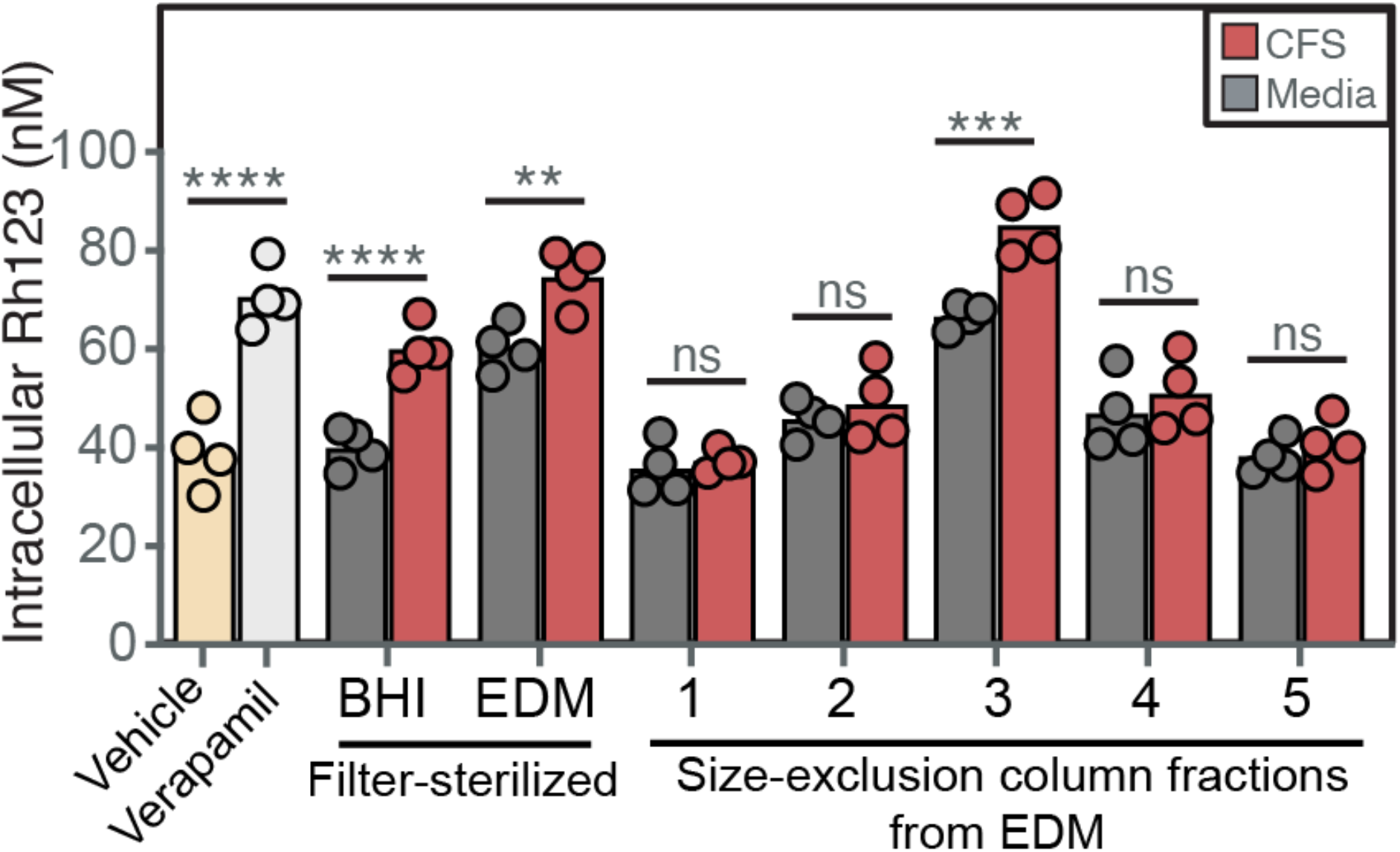
related to Figure 5. P-gp inhibitory activity is detected in *E. lenta* CFS cultured in EDM. *E. lenta* DSM2243 CFS from EDM cultures was processed through the activity-guided biochemical fractionation pipeline shown in **Fig. 5A** (n=4/group; two-way ANOVA with Tukey’s correction). Vehicle (0.02% DMSO) and 10 μM verapamil were included as controls. Filter-sterilized CFS cultured in EDM and BHI^+^ were included as controls for fractionations.

**Figure S3.**
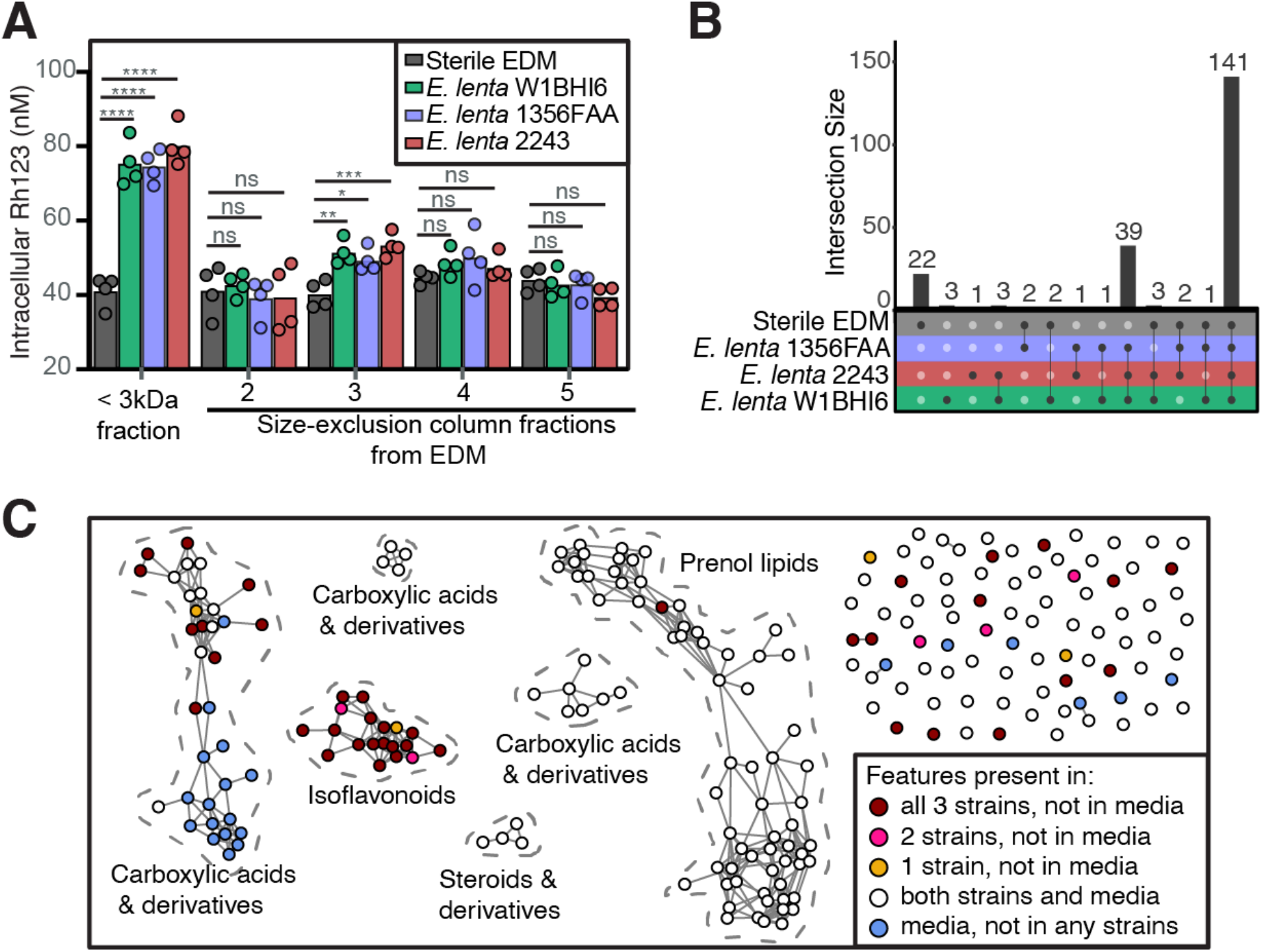
Expanded metabolomics screen links isoflavonoids with P-gp inhibition, related to Figure 6. CFS of 3 *E. lenta* strains (DSM2243, 1354FAA, W1BHI6) cultivated in EDM were processed through the activity-guided biochemical fractionation pipeline shown in **Fig. 5A** along with sterile EDM control. **(A)** Rh123 P-gp inhibition assay of each fraction (n=4/group; two-way ANOVA with Tukey’s correction). **(B)** Shared and unique features in the active fractions detected via untargeted metabolomics (see **Table S6**). **(C)** Functional annotations and network of features.

## REFERENCES

Alexander, Margaret, Qi Yan Ang, Renuka R. Nayak, Annamarie E. Bustion, Moriah Sandy, Bing Zhang, Vaibhav Upadhyay, Katherine S. Pollard, Susan V. Lynch, and Peter J. Turnbaugh. 2022. “Human Gut Bacterial Metabolism Drives Th17 Activation and Colitis.” Cell Host & Microbe 30 (1): 17–30.e9.

Altman, Russ B. 2007. “PharmGKB: A Logical Home for Knowledge Relating Genotype to Drug Response Phenotype.” Nature Genetics 39 (4): 426.

Bess, Elizabeth N., Jordan E. Bisanz, Fauna Yarza, Annamarie Bustion, Barry E. Rich, Xingnan Li, Seiya Kitamura, et al. 2020. “Genetic Basis for the Cooperative Bioactivation of Plant Lignans by Eggerthella Lenta and Other Human Gut Bacteria.” Nature Microbiology 5 (1): 56–66.

Bisanz, Jordan E., Paola Soto-Perez, Cecilia Noecker, Alexander A. Aksenov, Kathy N. Lam, Grace E. Kenney, Elizabeth N. Bess, et al. 2020. “A Genomic Toolkit for the Mechanistic Dissection of Intractable Human Gut Bacteria.” Cell Host & Microbe 27 (6): 1001–13.e9.

Bisanz, Jordan E., Peter Spanogiannopoulos, Lindsey M. Pieper, Annamarie E. Bustion, and Peter J. Turnbaugh. 2018. “How to Determine the Role of the Microbiome in Drug Disposition.” Drug Metabolism and Disposition: The Biological Fate of Chemicals 46 (11): 1588–95.

Brunton, Laurence, B. A. Chabner, and B. C. Knollmann. 2011. “Goodman and Gilman’s the Pharmacological Basis of Therapeutics. Twelfth.” New York, NY: McGraw-Hill.[Google Scholar]. http://www.clivar.org/sites/default/files/pdf-goodman-and-gilmans-the-pharmacological-basis-of-therapeutics-t-laurence-brunton-bruce-chabner-bjorn-knollman-pdf-download-free-book-7e42e5f.pdf.

Cheng, Alice G., Po-Yi Ho, Andrés Aranda-Díaz, Sunit Jain, Feiqiao B. Yu, Xiandong Meng, Min Wang, et al. 2022. “Design, Construction, and in Vivo Augmentation of a Complex Gut Microbiome.” Cell 185 (19): 3617–36.e19.

Cui, Jiahua, Xiaoyang Liu, and Larry M. C. Chow. 2019. “Flavonoids as P-Gp Inhibitors: A Systematic Review of SARs.” Current Medicinal Chemistry 26 (25): 4799–4831.

Dong, Xueyang, Ben Guthrie, Peter Turnbaugh, and Emily Balskus. 2022. “Genome Editing in an Actinobacterium Eggerthella Lenta.” In Preparation.

Duan, Zhenfeng, Edwin Choy, and Francis J. Hornicek. 2009. “NSC23925, Identified in a High-Throughput Cell-Based Screen, Reverses Multidrug Resistance.” PloS One 4 (10): e7415.

Elmeliegy, Mohamed, Manoli Vourvahis, Cen Guo, and Diane D. Wang. 2020. “Effect of P-Glycoprotein (P-Gp) Inducers on Exposure of P-Gp Substrates: Review of Clinical Drug-Drug Interaction Studies.” Clinical Pharmacokinetics 59 (6): 699–714.

Ernst, Madeleine, Kyo Bin Kang, Andrés Mauricio Caraballo-Rodríguez, Louis-Felix Nothias, Joe Wandy, Christopher Chen, Mingxun Wang, et al. 2019. “MolNetEnhancer: Enhanced Molecular Networks by Integrating Metabolome Mining and Annotation Tools.” Metabolites 9 (7). https://doi.org/10.3390/metabo9070144.

Foley, Sage E., Christine Tuohy, Merran Dunford, Michael J. Grey, Heidi De Luca, Caitlin Cawley, Rose L. Szabady, et al. 2021. “Gut Microbiota Regulation of P-Glycoprotein in the Intestinal Epithelium in Maintenance of Homeostasis.” Microbiome 9 (1): 183.

Fox, Elizabeth, and Susan E. Bates. 2007. “Tariquidar (XR9576): A P-Glycoprotein Drug Efflux Pump Inhibitor.” Expert Review of Anticancer Therapy 7 (4): 447–59.

Fu, Zidong Donna, Felcy P. Selwyn, Julia Yue Cui, and Curtis D. Klaassen. 2017. “RNA-Seq Profiling of Intestinal Expression of Xenobiotic Processing Genes in Germ-Free Mice.” Drug Metabolism and Disposition: The Biological Fate of Chemicals 45 (12): 1225–38.

Haiser, Henry J., David B. Gootenberg, Kelly Chatman, Gopal Sirasani, Emily P. Balskus, and Peter J. Turnbaugh. 2013. “Predicting and Manipulating Cardiac Drug Inactivation by the Human Gut Bacterium Eggerthella Lenta.” Science 341 (6143): 295–98.

Harwood, Caroline S., Gerhard Burchhardt, Heidrun Herrmann, and Georg Fuchs. 1998. “Anaerobic Metabolism of Aromatic Compounds via the Benzoyl-CoA Pathway.” FEMS Microbiology Reviews 22 (5): 439–58.

Hille, Russ. 1996. “The Mononuclear Molybdenum Enzymes.” Chemical Reviews 96 (7): 2757–2816.

Hooper, L. V., M. H. Wong, A. Thelin, L. Hansson, P. G. Falk, and J. I. Gordon. 2001. “Molecular Analysis of Commensal Host-Microbial Relationships in the Intestine.” Science 291 (5505): 881–84.

Javdan, Bahar, Jaime G. Lopez, Pranatchareeya Chankhamjon, Ying-Chiang J. Lee, Raphaella Hull, Qihao Wu, Xiaojuan Wang, Seema Chatterjee, and Mohamed S. Donia. 2020. “Personalized Mapping of Drug Metabolism by the Human Gut Microbiome.” Cell 181 (7): 1661–79.e22.

Katajamaa, Mikko, Jarkko Miettinen, and Matej Oresic. 2006. “MZmine: Toolbox for Processing and Visualization of Mass Spectrometry Based Molecular Profile Data.” Bioinformatics 22 (5): 634–36.

Koppel, Nitzan, Jordan E. Bisanz, Maria-Eirini Pandelia, Peter J. Turnbaugh, and Emily P. Balskus. 2018. “Discovery and Characterization of a Prevalent Human Gut Bacterial Enzyme Sufficient for the Inactivation of a Family of Plant Toxins.” eLife 7 (May). https://doi.org/10.7554/eLife.33953.

Kyaw, Than S., and Peter J. Turnbaugh. 2022. “Tiny Gatekeepers: Microbial Control of Host Drug Transporters.” Clinical Pharmacology and Therapeutics, June. https://doi.org/10.1002/cpt.2647.

Lam, Kathy N., Margaret Alexander, and Peter J. Turnbaugh. 2019. “Precision Medicine Goes Microscopic: Engineering the Microbiome to Improve Drug Outcomes.” Cell Host & Microbe 26 (1): 22–34.

Larsson, Erik, Valentina Tremaroli, Ying Shiuan Lee, Omry Koren, Intawat Nookaew, Ashwana Fricker, Jens Nielsen, Ruth E. Ley, and Fredrik Bäckhed. 2012. “Analysis of Gut Microbial Regulation of Host Gene Expression along the Length of the Gut and Regulation of Gut Microbial Ecology through MyD88.” Gut. https://doi.org/10.1136/gutjnl-2011-301104.

Ledwitch, Kaitlyn V., Morgan E. Gibbs, Robert W. Barnes, and Arthur G. Roberts. 2016. “Cooperativity between Verapamil and ATP Bound to the Efflux Transporter P-Glycoprotein.” Biochemical Pharmacology 118 (October): 96–108.

Limtrakul, P., O. Khantamat, and K. Pintha. 2005. “Inhibition of P-Glycoprotein Function and Expression by Kaempferol and Quercetin.” Journal of Chemotherapy 17 (1): 86–95.

Maini Rekdal Vayu, Elizabeth N. Bess, Jordan E. Bisanz, Peter J. Turnbaugh, and Emily P. Balskus. 2019. “Discovery and Inhibition of an Interspecies Gut Bacterial Pathway for Levodopa Metabolism.” Science 364 (6445). https://doi.org/10.1126/science.aau6323.

Maini Rekdal Vayu, Paola Nol Bernadino, Michael U. Luescher, Sina Kiamehr, Chip Le, Jordan E. Bisanz, Peter J. Turnbaugh, Elizabeth N. Bess, and Emily P. Balskus. 2020. “A Widely Distributed Metalloenzyme Class Enables Gut Microbial Metabolism of Host- and Diet-Derived Catechols.” eLife 9 (February). https://doi.org/10.7554/eLife.50845.

Mohimani, Hosein, Alexey Gurevich, Alexander Shlemov, Alla Mikheenko, Anton Korobeynikov, Liu Cao, Egor Shcherbin, Louis-Felix Nothias, Pieter C. Dorrestein, and Pavel A. Pevzner. 2018. “Dereplication of Microbial Metabolites through Database Search of Mass Spectra.” Nature Communications 9 (1): 4035.

Moore, Bradley S., Christian Hertweck, Jörn N. Hopke, Miho Izumikawa, John A. Kalaitzis, George Nilsen, Thomas O’Hare, et al. 2002. “Plant-like Biosynthetic Pathways in Bacteria: From Benzoic Acid to Chalcone.” Journal of Natural Products 65 (12): 1956–62.

Noecker, Cecilia, Juan Sanchez, Jordan E. Bisanz, Veronica Escalante, Margaret Alexander, Kai Trepka, Almut Heinken, et al. 2022. “Systems Biology Illuminates Alternative Metabolic Niches in the Human Gut Microbiome.” bioRxiv. https://doi.org/10.1101/2022.09.19.508335.

Nothias, Louis-Félix, Daniel Petras, Robin Schmid, Kai Dührkop, Johannes Rainer, Abinesh Sarvepalli, Ivan Protsyuk, et al. 2020. “Feature-Based Molecular Networking in the GNPS Analysis Environment.” Nature Methods 17 (9): 905–8.

Palmeira, A., E. Sousa, M. H. Vasconcelos, and M. M. Pinto. 2012. “Three Decades of P-Gp Inhibitors: Skimming through Several Generations and Scaffolds.” Current Medicinal Chemistry 19 (13): 1946–2025.

Pang, Zhiqiang, Jasmine Chong, Guangyan Zhou, David Anderson de Lima Morais, Le Chang, Michel Barrette, Carol Gauthier, Pierre-Étienne Jacques, Shuzhao Li, and Jianguo Xia. 2021. “MetaboAnalyst 5.0: Narrowing the Gap between Raw Spectra and Functional Insights.” Nucleic Acids Research 49 (W1): W388–96.

Pluskal, Tomás, Sandra Castillo, Alejandro Villar-Briones, and Matej Oresic. 2010. “MZmine 2: Modular Framework for Processing, Visualizing, and Analyzing Mass Spectrometry-Based Molecular Profile Data.” BMC Bioinformatics 11 (July): 395.

Rech, S., C. Wolin, and R. P. Gunsalus. 1996. “Properties of the Periplasmic ModA Molybdate-Binding Protein of Escherichia Coli.” The Journal of Biological Chemistry 271 (5): 2557–62.

Saha, Saha, V. Butler, H. Neu, and J. Lindenbaum. 1983. “Digoxin-Inactivating Bacteria: Identification in Human Gut Flora.” Science 220 (4594): 325–27.

Saksena, Seema, Sonia Goyal, Geetu Raheja, Varsha Singh, Maria Akhtar, Talat M. Nazir, Waddah A. Alrefai, Ravinder K. Gill, and Pradeep K. Dudeja. 2011. “Upregulation of P-Glycoprotein by Probiotics in Intestinal Epithelial Cells and in the Dextran Sulfate Sodium Model of Colitis in Mice.” American Journal of Physiology. Gastrointestinal and Liver Physiology 300 (6): G1115–23.

Siccardi, Dario, Karen L. Mumy, Daniel M. Wall, Jeffrey D. Bien, and Beth A. McCormick. 2008. “Salmonella Enterica Serovar Typhimurium Modulates P-Glycoprotein in the Intestinal Epithelium.” American Journal of Physiology. Gastrointestinal and Liver Physiology 294 (6): G1392–1400.

Silva, Ricardo R. da, Mingxun Wang, Louis-Félix Nothias, Justin J. J. van der Hooft, Andrés Mauricio Caraballo-Rodríguez, Evan Fox, Marcy J. Balunas, Jonathan L. Klassen, Norberto Peporine Lopes, and Pieter C. Dorrestein. 2018. “Propagating Annotations of Molecular Networks Using in Silico Fragmentation.” PLoS Computational Biology 14 (4): e1006089.

Spanogiannopoulos, Peter, Elizabeth N. Bess, Rachel N. Carmody, and Peter J. Turnbaugh. 2016. “The Microbial Pharmacists within Us: A Metagenomic View of Xenobiotic Metabolism.” Nature Reviews. Microbiology 14 (5): 273–87.

Sperry, J. F., and T. D. Wilkins. 1976. “Arginine, a Growth-Limiting Factor for Eubacterium Lentum.” Journal of Bacteriology 127 (2): 780–84.

Sugimoto, Yuki, Francine R. Camacho, Shuo Wang, Pranatchareeya Chankhamjon, Arman Odabas, Abhishek Biswas, Philip D. Jeffrey, and Mohamed S. Donia. 2019. “A Metagenomic Strategy for Harnessing the Chemical Repertoire of the Human Microbiome.” Science 366 (6471). https://doi.org/10.1126/science.aax9176.

Turnbaugh, Peter J., Vanessa K. Ridaura, Jeremiah J. Faith, Federico E. Rey, Rob Knight, and Jeffrey I. Gordon. 2009. “The Effect of Diet on the Human Gut Microbiome: A Metagenomic Analysis in Humanized Gnotobiotic Mice.” Science Translational Medicine 1 (6): 6ra14.

Wandy, Joe, Yunfeng Zhu, Justin J. J. van der Hooft, Rónán Daly, Michael P. Barrett, and Simon Rogers. 2018. “Ms2lda.org: Web-Based Topic Modelling for Substructure Discovery in Mass Spectrometry.” Bioinformatics 34 (2): 317–18.

Wang, Mingxun, Jeremy J. Carver, Vanessa V. Phelan, Laura M. Sanchez, Neha Garg, Yao Peng, Don Duy Nguyen, et al. 2016. “Sharing and Community Curation of Mass Spectrometry Data with Global Natural Products Social Molecular Networking.” Nature Biotechnology 34 (8): 828–37.

Wang, Yuhua, Yanlong Liu, Anju Sidhu, Zhenhua Ma, Craig McClain, and Wenke Feng. 2012. “Lactobacillus Rhamnosus GG Culture Supernatant Ameliorates Acute Alcohol-Induced Intestinal Permeability and Liver Injury.” American Journal of Physiology. Gastrointestinal and Liver Physiology 303 (1): G32–41.

Wishart, David S., Yannick D. Feunang, An C. Guo, Elvis J. Lo, Ana Marcu, Jason R. Grant, Tanvir Sajed, et al. 2018. “DrugBank 5.0: A Major Update to the DrugBank Database for 2018.” Nucleic Acids Research 46 (D1): D1074–82.

Yang, X., P. Yang, J. Shen, E. Osaka, E. Choy, G. Cote, D. Harmon, et al. 2014. “Prevention of Multidrug Resistance (MDR) in Osteosarcoma by NSC23925.” British Journal of Cancer 110 (12): 2896–2904.

Zhong, Qifeng, Bostjan Kobe, and Ulrike Kappler. 2020. “Molybdenum Enzymes and How They Support Virulence in Pathogenic Bacteria.” Frontiers in Microbiology 11 (December): 615860.

Zimmermann, Michael, Maria Zimmermann-Kogadeeva, Rebekka Wegmann, and Andrew L. Goodman. 2019. “Mapping Human Microbiome Drug Metabolism by Gut Bacteria and Their Genes.” Nature, June. https://doi.org/10.1038/s41586-019-1291-3.

Zou, Ling, Peter Spanogiannopoulos, Lindsey M. Pieper, Huan-Chieh Chien, Wenlong Cai, Natalia Khuri, Joshua Pottel, et al. 2020. “Bacterial Metabolism Rescues the Inhibition of Intestinal Drug Absorption by Food and Drug Additives.” Proceedings of the National Academy of Sciences 117 (27): 16009–18.

